# Distinct control of PERIOD2 degradation and circadian rhythms by the oncoprotein MDM2

**DOI:** 10.1101/286708

**Authors:** JingJing Liu, Xianlin Zou, Tetsuya Gotoh, Anne M. Brown, Liang Jiang, Jae Kyoung Kim, Carla V. Finkielstein

## Abstract

The circadian clock relies on post-translational modifications to set the timing for degradation of core regulatory components and, thus, sets clock progression. Ubiquitin-modifying enzymes targeting clock components for degradation are known to mostly recognize phosphorylated substrates. A case in point is the circadian factor *<u>PER</u>*IOD *<u>2</u>* (PER2) whose phospho-specific turnover involves its recognition by β-transducin repeat containing proteins (β-TrCPs). Yet, the existence of this unique mode of regulation of PER2’s stability falls short of explaining persistent oscillatory phenotypes reported in biological systems lacking functional elements of the phospho-dependent PER2 degradation machinery.

In this study, we challenge the phosphorylation-centric view that PER2 degradation enhances circadian rhythm robustness by *i*) identifying the PER2:MDM2 endogenous complex, *ii*) establishing PER2 as a previously uncharacterized substrate for MDM2, *iii*) revealing an alternative phosphorylation-independent mechanism for PER2 ubiquitin-mediated degradation, *iv*) pinpointing residues for ubiquitin modification, and *v*) establishing the importance of MDM2-mediated PER2 turnover for defining the circadian period length. Our results not only expand MDM2’s suite of specific substrates beyond the cell cycle to include circadian components but also uncover novel regulatory players that likely impact our view of how other mechanisms crosstalk and modulate the clock itself.

## INTRODUCTION

Circadian rhythms are endogenously-generated 24-h oscillations of biochemical, physiological, and behavioral processes that allow organisms to adapt to external environmental conditions. The coupling of the mammalian circadian system to various cellular process provides a means to understand the timing of when events take place in normal proliferative cells, a phenomenon believed to reflect evolutionary adaptation [for review see (1)]. As a result, identifying shared regulatory elements that operate under normal conditions and are relevant to the timely execution of cellular events might help establish nodes whose deregulation would be relevant to the understanding of human pathologies.

From a molecular standpoint, the circadian clock is formed by a transcriptional-translational feedback loop where expression of the core components drives the different phases of the daily cycle and whose protein products influence the cell’s biochemistry (2). In mammalian cells, the positive limb of the clock is driven by the heterodimer formed by the *<u>c</u>*ircadian *l*ocomotor *<u>o</u>*utput *<u>c</u>*ycles *<u>k</u>*aput (CLOCK) and the *<u>b</u>*rain and *<u>m</u>*uscle *<u>A</u>*rnt-*l*ike protein-*<u>1</u>* (BMAL1) complex, which initiates the transcription of *PERIOD* and *CRYTOCHROME* genes (*PER 1,2,3* and *CRY 1,2*) as well as other *<u>c</u>*lock-*<u>c</u>*ontrolled *<u>g</u>*enes (*ccgs*). Of note, several *ccgs* encode for cell cycle regulators. Dimerization of PER and CRY is relevant to the negative limb of the feedback loop as nuclear translocation of the complex further inhibits CLOCK/BMAL1 transcriptional activity [for review see (3)]. Thus, the stability of PER and CRY proteins is pertinent to the timing at which the termination of the repression phase takes place and the initiation of a new round of transcription begins. This process is mediated by distinct phosphorylation events in PER and CRY that precede E3-ligase-mediated ubiquitination and proteasomal degradation [for review see (3) and references within].

PERIOD 2 is a large protein with a well-defined N-terminus domain that is responsible for multiple protein-protein interactions, including homo- and hetero-dimerization among PER proteins (4). In addition, PER2 exhibits motifs and domains that play critical functional roles in its cellular localization (nuclear localization and export signal motifs), stability (binding domain for E3 ligase), and post-translationally-targeted modifications (including *<u>c</u>*asein *<u>k</u>*inase *<u>1</u>* ε/δ, CK1 ε/δ and *<u>g</u>*lycogen *<u>s</u>*ynthase *<u>k</u>*inase 3β, GSK3β, phosphorylation sites), which influence the periodic accumulation and distribution of PER2 in the cell [for review see (5)]. Furthermore, the stability of PER2, which seems to depend on its phosphorylation status (6, 7), is influenced by environmental stimuli and homeostatic cellular conditions (8–10) and is a critical determinant of the period length and phase of circadian rhythms (6,7,11). As a result, PER2 acts as a cellular rheostat that integrates signals and helps to robustly compensate for profound changes in environmental conditions that would otherwise affect the circadian clock.

Phosphorylation of PER2 by CK1 ε/δ can either stabilize or destabilize the circadian factor depending on what cluster site in PER2 is modified (9,11,12). Accordingly, PER2^S662G^, a PER2 variant linked to familial advanced sleep phase syndrome (13), contains a missense mutation that prevents priming-dependent phosphorylation of flanking sites by CK1 ε/δ, stabilizing PER2 independent of its cellular location (14). Conversely, a priming-independent cluster located in the C-terminus of PER2’s PAS domain is targeted by CK1 ε/δ and is required for ubiquitin ligase-mediated degradation of PER2 (15). Presently, our understanding of the molecular players involved in PER2 degradation is reduced to the sole role of β-TrCP, an F-box/WD40 repeat-containing substrate recognition subunit of the ubiquitin ligase complex SCF (*<u>S</u>*kp1-*<u>C</u>*ul1-*<u>F</u>*-box), that channels phosphorylation-dependent degradation of proteins (15, 16). The mammalian β-TrCP E3 ligase subfamily includes β-TrCP1 and β-TrCP2, both of which are closely related in sequence and indistinguishable in function but encoded by different genes (17). Biochemical evidence points to direct interactions between β-TrCP1/2 and PER1, but β-TrCP1, appears to be the sole form implicated in the binding of PER2 *in vitro* (15, 18). Regardless of these findings, there is no clear answer as to whether β-TrCP-targeted selectivity actually happens *in vivo* even though β-TrCP-mediated degradation contributes to generating cyclic levels of PER proteins relevant to the function of the clock (16). As has been noted, endogenous β-TrCPs’ activities depend on their localization and abundance in cells with β-TrCP1 being predominantly located in the nucleus and β-TrCP2, the most unstable form of both E3-ligases, being predominantly located in the cytoplasmic compartment (17).

Interestingly, findings show that overexpression of both dominant-negative forms of β-TrCP in cells neither increased PER2 stability nor accumulated phosphorylated PER2; instead, it resulted in rapid degradation of PER2 by a yet unknown mechanism (16). Similarly, expression of the dominant-negative form of CK1ε in a CK1δ^-/-^ background perturbs, but does not abrogate, circadian rhythms (19), a result that mimics one obtained using pharmacological inhibitors (20). More recently, Zhou and Kim *et al.*, have defined a phosphoswitch in PER2 that generates a three-stage kinetic degradation process (9). This mechanism allows fine-tuning of the stable fraction of PER2 and, therefore, adjusts the length of the circadian period due to diverse environmental stimuli (9). Remarkably, whereas the rapid initial decay in PER2 levels is phosphorylation-dependent and mediated by the activity of β-TrCP, the second “plateau” stage results from priming phosphorylation and accumulation of PER2 (9). Triggering PER2’s degradation in the third kinetic stage and during the falling phase of the circadian cycle are predicted to be independent of both phosphorylation and β-TrCP activity (9). More recently, findings using mice bearing loss-of-function mutations in β-TrCP1 and β-TrCP2 genes show that their behavioral wake/sleep phenotype is molecularly linked to PER degradation (21). Interestingly, the data show that PER proteins *“were still* [*ubiquitinated and*] *degraded, albeit at a slower rate* (21)*”* in double knockout β-TrCP1/2 cells. Thus, other E3 ligase(s) in addition to β-TrCP likely exist and contribute to PER’s overall stability.

We previously reported that PER2 forms a stable complex with the checkpoint and tumor suppressor protein p53 (22). The PER2:p53 complex undergoes time-of-day dependent nuclear-cytoplasmic shuttling, thus, generating an asymmetric distribution of each protein in different cellular compartments (23). In unstressed cells, PER2 mediates p53’s stability by binding p53’s C-terminus and preventing p53’s ubiquitination of targeted sites by the RING finger-containing E3 ligase *<u>m</u>*ouse *<u>d</u>*ouble *<u>m</u>*inute *<u>2</u>* homolog (MDM2) (22). Remarkably, PER2:p53:MDM2 co-exist as a trimeric and stable complex in the nuclear compartment, although p53 is released from the complex to become transcriptionally active once cells experience a genotoxic stimuli (24). As a result, we asked whether PER2 could also act as a *bona fide* substrate for the E3 ligase activity of MDM2 in the absence of p53. Unlike β-TrCP, MDM2 acts as a scaffold protein to facilitate catalysis by bringing the E2 ubiquitin-conjugating enzyme and substrate together in a phosphorylation-independent manner (17, 25). Our findings show that PER2 directly interacts with MDM2 opposite to the MDM2’s E3-RING domain and downstream of its p53-binding site. As a result, PER2 is efficiently ubiquitinated *in vitro* and in cells at numerous sites by MDM2 in a process that is preferentially mediated by UbcH5a, a robust E2-ubiquitin conjugating enzyme with innate preference for various polyubiquitin chain linkages (26). Furthermore, MDM2-mediated ubiquitination on PER2 is independent of phosphorylation. Accordingly, PER2’s half-life is critically modulated by MDM2’s levels and its enzymatic activity as shown in cells where MDM2’s expression is either enhanced or silenced and its catalytic activity pharmacologically inhibited. Consequently, direct manipulation of MDM2 expression influences period length by acting on PER2’s stability. Therefore, our results provide evidence to advocate for tight control of PER2’s turnover in cells that expand the phosphorylation-centric view of its degradation.

## MATERIALS AND METHODS

### Plasmid constructs

The human PER2 (NM_022817), p53 (NM_000546), Mdm2 (NM_002392), β-TrCP1 (NP_003930), and CK1ε (BC006490, Addgene) full-length cDNAs were cloned downstream from a tag encoding sequence into pCS2+3xFLAG-, and *(myc)_6_*-tag vectors (FLAG-PER2, FLAG-p53, FLAG-MDM2, FLAG-β-TrCP, FLAG-CK1ε, *myc*-PER2, *myc*-p53, *myc*-Mdm2) modified for ligation-independent cloning (LIC, Novagen). Various mutants of MDM2 (*e.g.*, MDM2C^470^A) and PER2 (*e.g.*, PER2S^662^A) were generated from the FLAG- and *myc*-tagged templates, respectively, using QuikChange II site-directed mutagenesis and following manufacturer’s instructions (Agilent). Deletion constructs of MDM2 [MDM2(1-117), MDM2(117-497), MDM2(1-230), MDM2(230-497), MDM2(1-434), MDM2(434-497)], p53 [p53Δ30, comprises residues 1-363 in p53], and PER2 [PER2(1-682), PER2(356-872), PER2(683-872), PER2(873-1,255)] were obtained by PCR amplification and subcloning in either pCS2+3xFLAG- or *(myc)_6_*-tag vectors. Various lengths of cDNA fragments of PER2 were cloned into the *SalI*/*NotI* sites of pGEX-4T-3. Fragments of PER2 comprising residues 1-172, 173-355, 356-574, 575-682, 683-872, 873-1,120, and 1,121-1,255 are referred in the text as: GST-PER2(1-172), GST-PER2(173-355), GST-PER2(356-574),GST-PER2(575-682), GST-PER2(683-872), GST-PER2(873-1120), GST-PER2(1121-1255), respectively.

### Bacterial two-hybrid screening

The two-hybrid interaction screening was performed using the BacterioMatch II system (Stratagene) following manufacturer’s instructions. A specific bait (pBT-*PER2*) and target plasmid pair from a liver library (pTRG cDNA library) were co-transformed with the bait vector plus the pTRG target vector. Selection was performed as indicated in (22) and positive colonies were transferred from selective screening medium onto a dual selective screening medium plate containing 3-amino-1,2,4-triazole (3-AT) and streptomycin. The pBT-*LFG2*/pTRG-*Gal11*^P^ co-transformant was used as a positive control whereas co-transformation of pBT-*PER2* with either empty pTRG or pTRG-*Gal11^P^* vectors were used as negative controls. All positive cDNA clones were isolated and sequenced, with some of them already having been reported, and their interaction functionally verified (22).

### Cell culture, transient transfections, and treatments

Human colorectal carcinoma HCT116 [*TP53*(+/+), *PER2*(+/+)] and human non-small cell lung carcinoma H1299 cell lines were purchased from the American Type Culture Collection (ATCC) and propagated according to manufacturer’s recommendations. The H1299 cells contained a homozygous partial deletion of the *TP53* gene that results in the absence of p53 expression. The HCT116 null-isogenic clone [*TP53*(-/-), *PER2*(+/+)] was purchased from GRCF Biorepository and Cell Center (Johns Hopkins School of Medicine) and maintained in McCoy’s 5A modified medium containing 10% Fetal Bovine Serum (FBS), 50 U/ml of penicillin and 50 μg/ml of streptomycin. The MEF^mPer2::LUC^ cells (kind gift of S. Kojima, Virginia Tech) were cultured in Dulbecco’s Modified Eagle’s Medium (DMEM, 4.5 g/l glucose) supplemented with 10% fetal bovine serum (FBS), 50 U/ml of penicillin, and 50 μg/ml of streptomycin, and maintained at 37°C and 5% CO_2_.

Plasmid transfections were performed at 50-80% cell confluency and optimized using Lipofectamine LTX (Invitrogen) and HyClone HyQ-RS reduced serum medium (GE Healthcare) following manufacturer’s instructions. Proteins were allowed to express for several hours before cells were either harvested or circadian synchronized. Synchronization was by serum shock (27) or dexamethasone treatment (28). Lysates were from cells collected at the indicated times, with t=0 occurring just prior to cycloheximide (CHX, 100 μg/ml) addition.

For siRNA transfections, HCT116 p53^+/+^ cells were grown in McCoy’s 5A media containing 10% FBS, penicillin (50 U/ml), and streptomycin (50 μg/ml) until reaching 60-80% confluency. Knockdown was optimized using Dharmafect 2 reagent (GE Dharmacon) to deliver siRNAs targeting either MDM2 (5’-GAGATTTGTTTGGCGTGCCAAGCTT-3’) or β-TrCP1 (5’-CGGAAACTCTCAGCAAGCTATGAAA-3’) following manufacturer’s instructions. A scramble siRNA sequence with no homology to any known mammalian gene, served as control. Forty-eight hours after transfection, cells were serum shocked for 2 h after which the media was replaced with a serum-free version and cycloheximide was added. Samples were collected at different times after treatment and extracts were prepared in NP-40 lysis buffer containing 10 mM Tris-HCl (pH 7.5), 137 mM NaCl, 1mM EDTA, 10% glycerol, 0.5% NP-40, 80 mM β-glycerophosphate, 1mM Na_3_VO_4_, 10 mM NaF, and protease inhibitors (10 μM leupeptin, 1 μM aprotinin A, and 0.4 μM pepstatin).

Lastly, endogenous levels of PER2 were monitored in HCT116 cells treated with CHX and incubated with sempervirine nitrate (named SN, 1μg/ml, ChromaDex Inc.), PF670462 (named PF670, 0.1μM or 1μM, Cayman Chemical Co.), or a combination of both inhibitors throughout the time course analyzed. The vehicle (DMSO) was used as control.

### Immunoprecipitation and immunoblot assays

Immunoprecipitation of protein complexes were from either transfected cell extracts or *in vitro* binding reactions. Unless indicated, proteins were in NP-40 lysis buffer, and extracts (0.5-1 mg) were incubated by rotation for either α-FLAG M2 agarose beads (Sigma-Aldrich) or α-*myc* (9E10) beads (Santa Cruz Biotechnology) for either 2 h or overnight at 4°C, respectively. In other cases, immunoprecipitations were carried out in a two-step procedure with extracts first being incubated with an uncoupled antibody (α-FLAG, α-*myc*, or α-PER2) overnight at 4°C before the addition of protein A beads (50% slurry; Sigma-Aldrich). Bound beads were washed four times with wash buffer (20 mM Tris-HCl (pH 7.5), 100 mM NaCl, 5 mM EDTA, 0.1% Triton X-100, and 0.5 mM PMSF) before the addition of Laemmli buffer. Complexes were resolved by SDS-PAGE and immunoblotting using the specific antibodies indicated in each case. Primary antibodies were α-FLAG (Sigma-Aldrich), α-*myc* (Santa Cruz Biotechnology), α-PER2 (Sigma-Aldrich), α-MDM2 (Santa Cruz Biotechnology), α-β-TrCP1 (Cell Signaling Technology), α-p53 (DO1 clone, Santa Cruz Biotechnology), and α-ubiquitin (Enzo Life Sciences). Secondary antibodies were horseradish peroxidase-conjugated α-rabbit or α-mouse IgGs (Invitrogen) and chemiluminescence reactions were performed using the SuperSignal West Pico Substrate (Pierce).

### In vitro binding and epitope blocking assays

*In vitro* transcription and translation of either pCS2+*myc*- or –FLAG PER2, β-TrCP1, β-TrCP1ΔF, MDM2, MDM2(C^470^A), and p53 were carried out using the SP6 high-yield TNT system (Promega) following manufacturer’s instructions. As indicated in each case, aliquots (1-4μl) of the indicated recombinant proteins were pre-incubated for 15 min at room temperature to allow complex formation before adding NP-40 lysis buffer. Epitope blocking was performed by pre-incubating *in vitro* the transcribed and translated FLAG-MDM2(C^470^A) with α-4B11, -4B2, or –SMP14 antibody (0.1 mg/ml each, Calbiochem) for 2 h at 4°C before adding recombinant *myc*-PER2. Binding reactions were allowed to proceed overnight at 4°C with rotation. In other experiments, binding of *myc*-MDM2 or - MDM2(C^470^A) proteins was evaluated in λPPase [200U (New England Biolabs), 15 min at 25 °C]-treated recombinant FLAG-PER2 samples. Reactions were diluted in NP-40 lysis buffer and complexes were immunoprecipitated using α-FLAG antibody (Sigma) and protein A beads (50% slurry) as described earlier.

### Protein pull-down assay

Recombinant GST-tagged PER2 proteins were expressed in *E. coli* strain *Rosetta* (Novagen) and purified using glutathione sepharose affinity chromatography following manufacturer’s instructions (GE Healthcare). Pull-down experiments were carried out using 5 μg of each recombinant GST-tagged protein-bound beads, or an equivalent amount of GST bound glutathione beads as control, and 4 μl of *in vitro* transcribed and translated [^35^S]-FLAG-MDM2 in binding buffer containing 20 mM Tris-HCl (pH 7.4), 100 mM NaCl, 5 mM EDTA, and 0.1% Triton X-100. Reactions were incubated for 1 h at 4°C with rotation, after which, bead-bound complexes were washed with binding buffer containing either low (100 mM) or high (1M) salt concentration. Bound proteins were analyzed by SDS-PAGE and autoradiography.

### Ubiquitination and degradation assays

Aliquots (1-4 μl) of *in vitro* transcribed and translated tagged-proteins [FLAG-, *myc*-, or *myc*-DO (DO epitope sequence is EPPLSQETFSDLWKL)], or a combination of them, were allowed to bind before adding 1x ubiquitination buffer (Enzo Life Sciences), 1 mM dithiothreitol, 20 μg/ml ubiquitin-aldehyde, 600 μg/ml ubiquitin, 1x ATP-energy regeneration system (5 mM ATP/Mg^2+^; Enzo Life Sciences), 40 μM MG132 (Cayman Chemical Co.), and 1 mg/ml of HeLa S100 lysate fraction (Enzo Life Sciences) or 1xE1 (ubiquitin activating enzyme, Enzo Life Sciences) and 1x UbcH5a, b, or c (human ubiquitin conjugating enzyme, human Enzo Life Sciences) to a final volume of 10-15 μl. Reactions were then incubated for 1 h at 37°C in a water bath, except for Supplementary Figure S3B where the incubation time varied as indicated in the figure. In other experiments, PER2 recombinant proteins were pre-treated with λPPase [200U (New England Biolabs), 15 min at 25 °C] before the ubiquitination reaction was carried out in the presence of tagged MDM2 as aforementioned. Following, Laemmli sample buffer was added and ubiquitinated proteins were either resolved by SDS-PAGE and detected by immunoblotting or immunoprecipitated following the two-step protocol described in the section above.

Detection of ubiquitinatinated forms of PER2 in cells was carried out by co-transfecting HCT116 [*TP53*(+/+), *PER2*(+/+)] cells with pCS2+FLAG-PER2 and either pCS2+*myc*-MDM2 or pCS2+*myc*-MDM2(C470A) plasmids. Cells were maintained in complete media for 24h to allow for the recombinant proteins’ expression before adding MG132 (50 μM) and ubiquitin aldehyde (5 nM). Cells were harvested 4 h later and lysates were immunoprecipitated using α-FLAG antibody as described. Proteins were resolved by SDS–PAGE and ubiquitinated forms of PER2 were detected by immunoblotting using an α-ubiquitin antibody.

### Homology model generation and protein-protein docking

The I-TASSER Server (29) was used to create homology models of PER2(683-872) wild-type and mutant variants. Sequences were uploaded to the server in FASTA format. There were no restraints guiding modeling, homologous templates were not excluded, and secondary structures for specific residues were unbiased. The server uses templates from the PDB database to predict secondary structure of the query protein using LOMETS 4 (Local Meta-Threading-Server). Alternatively, the server uses *ab initio* modeling to assign secondary structures. Clustering is then performed to find the lowest free-energy model using SPICKER (30). The model with the highest C-score value was then energy minimized using the Molecular Operating Environment utilizing Amber12EHT parameters and subsequently validated using online servers including SWISS-MODEL, ProSA, and Verify3D. The models of all three constructs showed reasonable energies relevant to their Anolea, Procheck, and z-scores, as well as favorable 3D structure and side chain placements, and were deemed acceptable. Herein, the three models were validated and used in confidence in further protein-ubiquitin docking experiments.

Protein binding interfaces were predicted by docking between ubiquitin [PDB:1UBQ (31)] and each model of the PER2 fragments. The Schrödinger software suite (2017.2) and the BioLuminate interface, which utilizes the PIPER docking module, were used for interface determination. No biased or interfaced residue was set at the onset of docking. All PER2 models were treated equally in regard to how and where ubiquitin molecules were predicted to interact. Thirty structures for each PER2:ubiquitin docking pair were obtained and clustered using the pairwise root mean square deviation (RMSD) and key residues located at the interface identified. Data files are available from the Virginia Tech Institutional Data Repository, VTechData, <u>doi:10.7294/W4JW8C2R</u> (DOI to be awarded at publication and deposit).

### Analysis of protein half-life

Accumulation and half-life of endogenous proteins in HCT116 cell extracts (20-80 μg) was monitored by immunoblotting samples collected at different times after CHX addition as indicated in the figure legends. Protein bands were quantified by immunoblot analysis using Bio-Rad ImageLab 5.1 software/Gel Doc XR+ system and values were normalized to tubulin levels. Unless indicated, the percentage of remaining protein was normalized to t=0 and the data fitted using Microsoft Excel.

In other experiments, the half-life of PER2 was measured in MEF^mPer2::LUC^ cells by luminescence recording. Seeded cells were synchronized with dexamethasone (100 nM, 2 h) and maintained in media containing phenol-red-free DMEM, 50 μM luciferin (Biosynth), 2% FBS, 1% penicillin/streptomycin, 1% L-glutamine (Invitrogen) in a lumicycle instrument (t= 0h). Addition of CHX (40 μg/ml) and DMSO (1% v/v), PF670462 (1μM), SN (1 μg/ml), or both inhibitors occurred during rising (t= 24h) or falling (t= 33h) phases. Three biological experiments were performed in parallel with each treatment being plated in triplicate. Data were normalized to the PER2::LUC signal from untreated cells. The PER2 half-life was determined at the time in which PER2::LUC signal was 50% of the initial detected amount as degradation curves were not exponential.

### Real-time bioluminescence assays

Cells, MEF^mPer2::LUC^, were seeded in 35 mm dishes and circadian synchronized by dexamethasone treatment (100 nM, 2 h). Following media replacement as described above, cells were allowed to stabilize in a LumiCycle 32-channel automated luminometer (Actimetrics) placed in a 37°C incubator for 24 h before sempervirine (1μg/ml), PF670462 (0.1 or 1 μM), or both inhibitors were added. In these assays, bioluminescence was continuously recorded for at least 5 additional days and data were analyzed using the LumiCycle analysis software (Actimetrics).

In other experiments, MEF^mPer2::LUC^ cells were transiently transfected with either pCS2+*myc*-*MDM2* or *siRNA MDM2* for 24 or 48 h, respectively, before dexamethasone synchronization. Following media exchange, bioluminiscence was recorded for at least 5 additional days. In each case, raw data was collected after dexamethasone synchronization (t=0) and for the remainder of the experiment. Raw data beginning t=24 h after synchronization was considered when calculating the circadian period length.

Period length was calculated using the Lumicycle data analysis software (Actimetrics). For all experiments, mean and errors were calculated based on at least triplicates.

## RESULTS

In vertebrates, phosphorylation-dependent β-TrCP-mediated ubiquitination and proteasomal degradation provides a means of regulating endogenous levels of PER2 in the cell and, thus, its daily accumulation [for review see (8)]. A large body of evidence supports a more central role for PER2 as the integrator of intracellular signals and as a sensor of environmental conditions. Thus, much effort has been devoted to understanding how various phosphorylation events determine PER2’s degradation rate (8-10,32,33) whereas other phosphorylation-independent mechanisms of degradation have remained largely unexplored. Building on our previous finding that PER2 forms a stable complex with p53 and MDM2 (22, 24), we evaluated whether the RING E3 ligase provides an alternative route for degradation of PER2 that is independent of phosphorylation and, at the same time, influences the circadian period.

### The oncogenic E3 ligase MDM2 interacts with PER2 in the absence of p53

Herein, and using a bacterial two-hybrid system, we report the identification of the human MDM2 homolog as a direct interactor of PER2 suggesting that, in addition to the already identified PER2:p53:MDM2 nuclear complex (24), PER2:MDM2 might exist as its own entity and that this association might be independent of p53 binding (Figure 1A). Consequently, we then evaluated the presence of the PER2:MDM2 complex in cells lacking endogenous expression of p53. Initial experiments were carried out using colorectal HCT116 cells [*TP53*(+/+), *PER2*(+/+); named HCT116^p53+/+^ hereafter] and, to avoid confounding variables, its null-isogenic HCT116 cell variant lacking p53 expression [*TP53*(-/-), *PER2*(+/+); named HCT116^p53-/-^ hereafter] (34). As p53 and MDM2 form a regulatory feedback loop in which p53 transcriptionally up-regulates MDM2 expression, cells lacking p53 protein usually exhibit constitutively low levels of MDM2 expression, which is enhanced by MDM2’s self-ubiquitination activity and increased turnover [for review see (35)]. To circumvent this problem, HCT116^p53-/-^ cells were transfected with *myc*-tagged MDM2 and PER2:MDM2 association was detected by immunoprecipitation of endogenous PER2 (Figure 1B). Accordingly, α-PER2, but not control IgG, antibody brings down PER2-associated MDM2 in both HCT116^p53+/+^ and HCT116^p53-/-^, further supporting their p53-independent interaction. Similar results were obtained using a human non-small cell lung carcinoma line (H1299) that possesses a homozygous partial deletion of the *TP53* gene. In this case, complexes were detected in cells co-transfected with *myc*-PER2 and FLAG-MDM2, its ubiquitin ligase activity-deficient mutant FLAG-MDM2(C^470^A) (36), or the E3 ligase β-TrCP1 (Supplementary Figure S1A). These results prompted us to map the region of binding between PER2 and MDM2 to better understand the interplay among these molecules.

**Figure 1.**
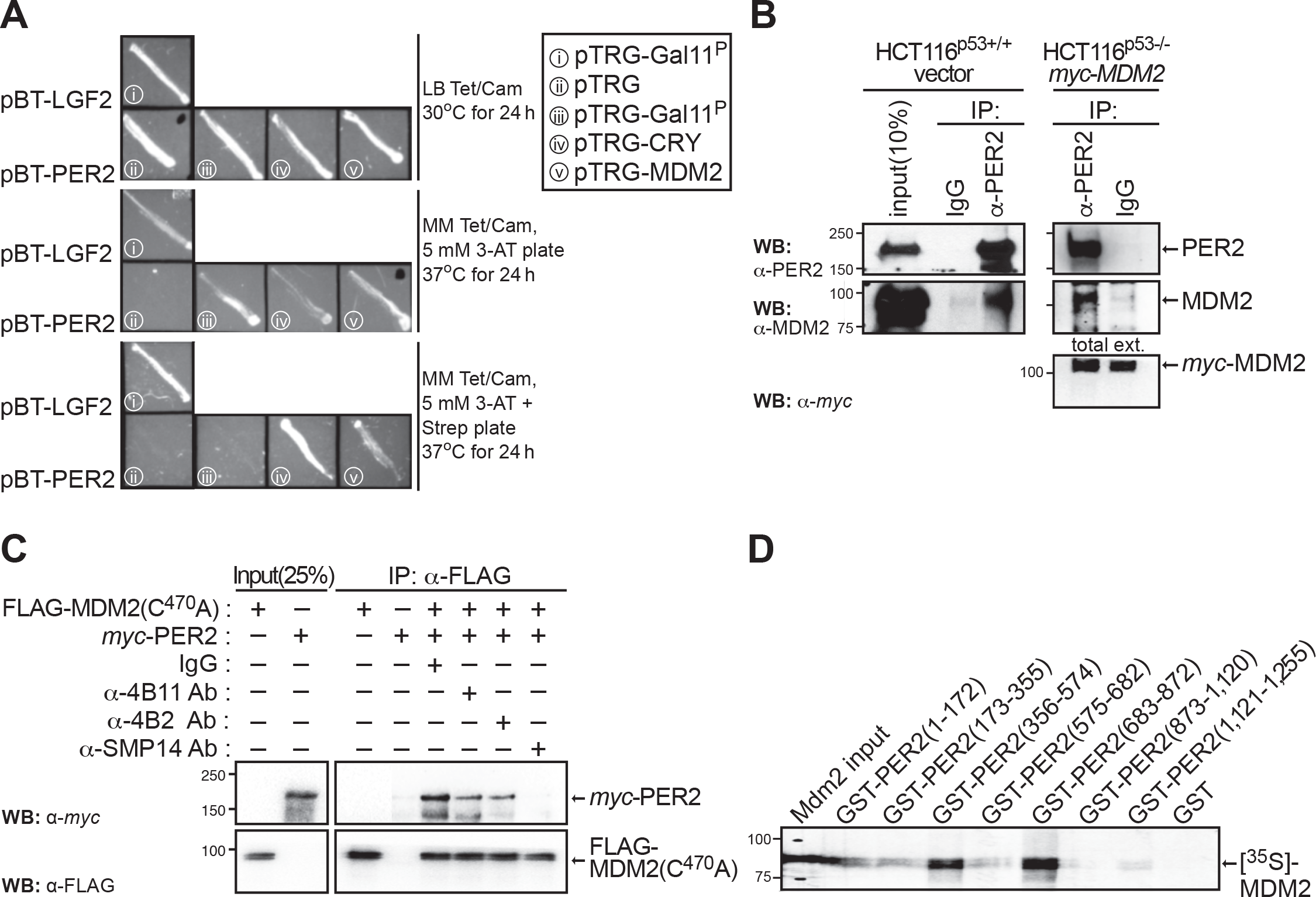
The circadian factor PERIOD 2 (PER2) directly interacts with MDM2. (**A**) A bacterial two-hybrid screening was developed to identify direct protein interactors of PER2 *in vivo* based on transcriptional activation. Positive clones encoding putative interacting proteins were maintained on LB tetracycline/chloramphenicol media (Tet/Cam, *upper panel*), and patched in His-dropout selective media containing either 3-amino-1,2,4-triazole (3-AT, *middle panel*) or 3-AT and streptomycin (Strep, *lower panel*). Patches of co-transformants served as positive (pBT-LFG2 and pTRG-Gal11^P^) and negative (pBT-PER2 and pTRG empty vector) controls. (**B**) Extracts from isogenic HCT116 cells (p53+/+ or p53-/-) were analyzed for the presence of endogenous PER2:MDM2 (*left panels*) or PER2:*myc*-MDM2 (*right panels*) complexes by immunoprecipitation using α-PER2 antibody and blotting using the indicated antibodies. IgG was used as the negative control. (**C**) Competition experiments were carried out by pre-incubating FLAG-MDM2(C^470^A) with each of the indicated α-MDM2 antibodies (α-4B11, -4B2, -SMP14) before adding recombinant *myc*-PER2. Protein binding was monitored in FLAG-bound beads by immunoblotting. IgG was used as negative control. (**D**) Pull-down assay was carried out using recombinant GST-tagged PER2 protein fragments comprising various lengths of PER2 [GST-PER2(1-172), GST-PER2(173-355), GST-PER2(356-574), GST-PER2(575-682), GST-PER2(683-872), GST-PER2(873-1,120), GST-PER2(1,121-1,255)], and radiolabeled [^35^S]-MDM2. The GST protein was used as a negative control. In all cases, molecular weight markers are indicated on the left (kDa). Immunoblot data from **B** and **C** were originated from a single experiment that was repeated three times with similar results. **D** was repeated twice.

Epitope mapping was carried out by pre-incubating recombinant FLAG-MDM2(C^470^A) with various epitope-specific antibodies that recognize native conformations in MDM2 [(37), 4B2: residues 19 to 59; SMP14: residues 154 to 167; 4B11: residues 383 to 491 (comprises the RING domain)] before adding recombinant *myc*-PER2 (Supplementary Figure S1B). As shown in Figure 1C, pre-incubation with the α-MDM2 clone SMP14 completely abolished PER2 binding, suggesting that the epitope comprising residues 154-167 in MDM2 is critical for the stability of the PER2:MDM2 interaction. Accordingly, immunoprecipitation of FLAG-MDM2 recombinant proteins engineered to include various functional domains confirmed that the N-terminus hydrophobic pocket in MDM2 (residues 1 to ∼110) is dispensable for PER2 recognition, even though contacts besides the SMP14 epitope exist between PER2 and MDM2 (Supplementary Figure S1C). Conversely, pull-down experiments using various recombinant GST-expressed fragments of PER2 [named GST-PER2 (1-172), GST-PER2 (173-355), GST-PER2 (356-574), GST-PER2 (575-682), GST-PER2 (683-872), GST-PER2 (873-1,120), GST-PER2 (1,121-1,255)] and [^35^S]-MDM2 showed that association primarily occurs at the C-terminus of the PER2 PAS domain, residues 356 to 574, and in a further inward region (residues 683 to 872) that is heavily post-translationally modified (Figure 1D). Overall, our findings establish the existence of a PER2:MDM2 complex whose association is independent of the presence of p53, suggesting that E3 ligases other than β-TrCP may be acting on PER2.

### Period 2 is a *bona fide* substrate of MDM2

As PER2 binds MDM2 opposite its RING domain, we asked whether MDM2’s catalytic activity could result in PER2 ubiquitination. To test this possibility, we evaluated PER2 ubiquitination in a cell-free system enriched in E1 and E2 enzymes containing *in vitro* transcribed and translated FLAG-MDM2 or FLAG-MDM2(C^470^A) and multi-tagged *myc*-DO-PER2 proteins (Figure 2A). Our data showed that ubiquitinated forms of PER2, depicted as a high molecular weight ladder, were distinguishable when the substrate was incubated in the presence of wild-type MDM2, but not its ligase-deficient mutant form (Figure 2A, lanes 2 *vs.* 3). Similarly, ubiquitinated forms of PER2 were detected in immunoprecipitated samples from *in vitro* reactions performed in the presence of FLAG-ubiquitin (Supplementary Figure S2A) and from lysates where co-transfected cells were maintained in the presence of the proteasome inhibitor MG132 (Supplementary Figure 2B).

**Figure 2.**
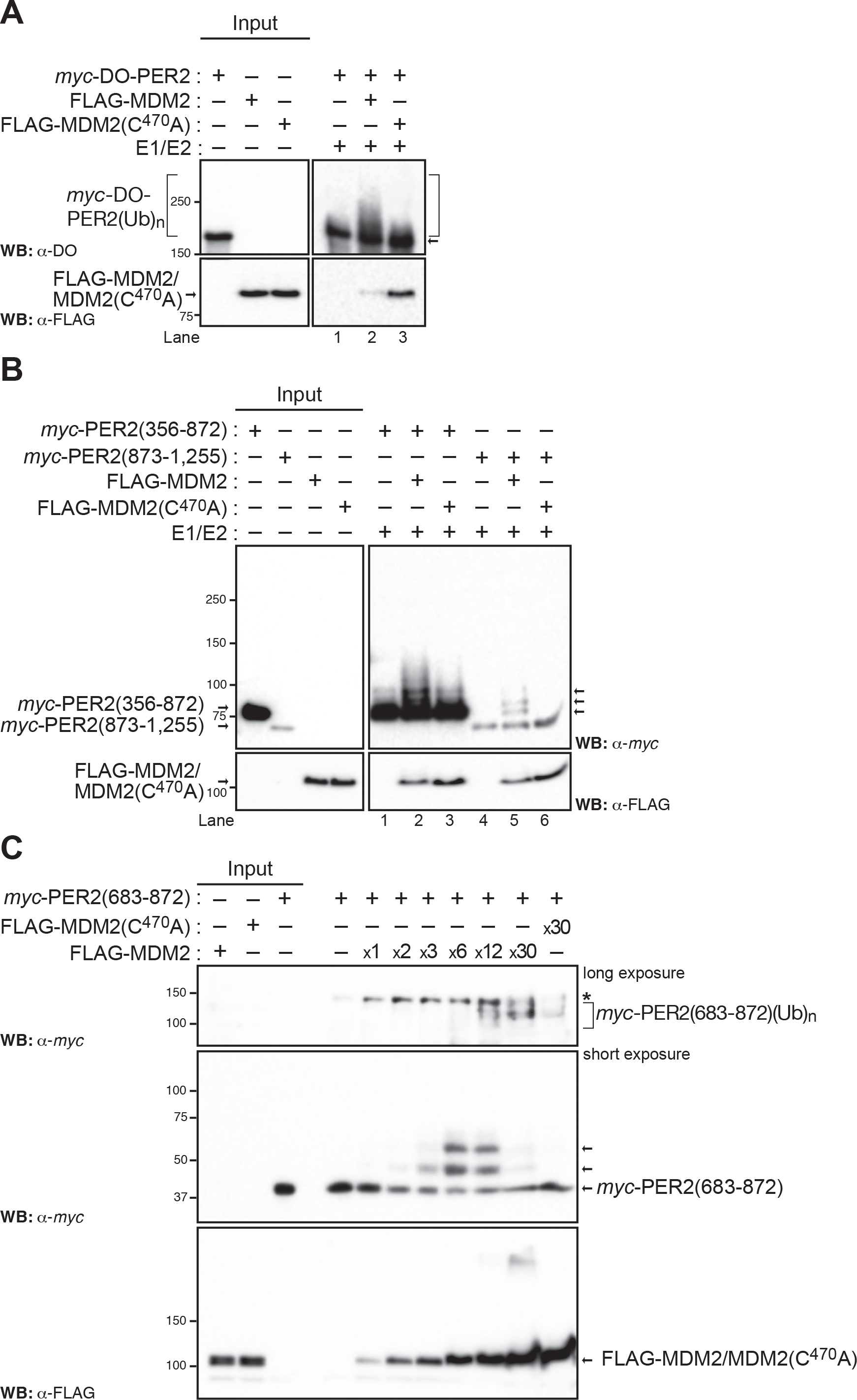
The E3-ligase MDM2 targets PER2 for ubiquitination. (A) *In vitro* ubiquitination reactions were carried out using recombinant transcribed and translated *myc*-DO-PER2, FLAG-tagged MDM2 or MDM2(C^470^A) proteins and E1/E2 enzymes as described in Materials and Methods and samples were analyzed by immunoblotting using α-tagged antibodies. (B) Fragments of PER2 comprising residues 356-872 and 873-1,255 were expressed as *myc*-tagged recombinant proteins, assessed for ubiquitination *in vitro*, and products detected by immunoblotting. (C) *In vitro* ubiquitination reactions were carried out using various enzyme:substrate ratios in a mix optimized as in Figure S3. Poly-ubiquitination forms of PER2(683-872) are indicated with a bracket (*upper panel*). Levels of E3 ligases are shown by blotting in the *lower panel*. In each case, immunoblot data was repeated three times with consistent results.

Next, we asked whether ubiquitin modifications in PER2 were confined to specific domains in the protein. We turned our attention to three relevant regions in PER2: *i*) the N-terminus PAS domain and its C-terminus extension (residues 1-682), which is responsible for PER2’s role as a transcriptional regulator, *ii*) a middle region, heavily post-translationally modified, largely involved in protein-protein interactions and in cellular shuttling of PER2 (residues 356-872), and *iii*) a C-terminus fragment (873-1,255), which is known to directly interact with various ligands and with PER2’s counterpart CRY (38,39) (Figure 2B and Supplementary Figure S2C). To this end, we reconstituted a functional E1/E2/MDM2 or MDM2(C^470^A) ubiquitin ligase *in vitro* system and used various recombinantly-expressed tagged fragments of PER2 as substrates. Results showed that PER2(1-682), PER2(356-872), and PER2(873-1,255) were able to incorporate multiple ubiquitin moieties when incubated with wild-type MDM2, but not MDM2(C^470^A), as it is depicted by the presence of a ladder in immunoblots (Figure 2B, lanes 2 *vs.* 3 and 5 *vs.* 6 and Supplementary Figure S2C, lane 2 *vs.* 3, *right arrows*).

We previously established that residues within the center region of PER2 lay at the interface of its interaction with p53 and facilitate the formation of a stable complex where circadian and checkpoint signals converge (22, 24). Thus, we turned our attention to defining ubiquitination events taking place, specifically, within residues 683 to 872 in PER2.

As shown for p53, whereas ubiquitination by MDM2 is influenced by various factors including the enzyme:substrate ratio, the incorporation of diverse ubiquitin chains in the substrate results from the multivalent nature of linkage-specific conjugations (40, 41). Therefore, we initially optimized the reaction conditions to ensure targeting of PER2(683-872) by MDM2 was taking place within the initial velocity region and the ubiquitin conjugation biochemically defined (Supplementary Figure S3). Initially, we tested scenarios in which the amount of ATP-Mg^2+^ chelate (Supplementary Figure 3A), ubiquitin co-substrate (Supplementary Figure S3A), and time-dependent accumulation of products were varied (Supplementary Figure S3B).

Next, as MDM2 functions with various E2 ubiquitin-conjugating enzymes from the UbcH5 family (a, b, and c) to mediate either specific ubiquitin linkages or more promiscuous ones in different substrates (41), we tested the relevance of UbcH5 specificity for MDM2-mediated ubiquitination of PER2(683-872) (Supplementary Figure S3C, *left panel*). *In vitro* ubiquitination reactions were performed using recombinant enzymes (E1 and UbcH5a-c) and tagged MDM2, MDM2(C^470^A), and PER2(683-872) substrates. The p53 protein was used as a positive control as it is efficiently modified by MDM2 in the presence of either ubiquitin conjugated UbcH5 enzyme (Supplementary Figure S3C, *right panel*). Results showed that all UbcH5 enzymes promoted the addition of, at least, a single ubiquitin molecule and that UbcH5a appeared to be more effective in catalyzing the incorporation of at least a second molecule (Supplementary Figure S3C, *left panel*). Among the eight possible ubiquitin linkages, UbcH5a displays selectivity for Lys^11^, Lys^48^, and Lys^63^ in promoting ubiquitin chain initiation (42). Of these, Lys^11^ and the canonical Lys^48^ linkages were involved in the formation of poly-ubiquitin chains and proteasome-mediated turnover whereas the Lys^63^ linkage plays a non-degradative role and is usually involved in protein recruitment and localization (42). Lastly, we confirmed a dose-dependent effect of MDM2 in the accumulation of slower-migrating poly-ubiquitinated forms of PER2(683-872) that were undetectable when the reaction took place in the presence of MDM2(C^470^A), (Figure 2C). Thus, our results allude to the existence of early ubiquitination events within PER2 from which poly-ubiquitination chains can be built upon.

### Binding of p53 to PER2 abrogates MDM2-mediated ubiquitination

To gain further insight into the role that MDM2 plays in PER2 function, we focused our efforts on identifying relevant Lys residues that could be targeted for modification. A highly conserved cluster of Lys residues (K^789^, K^790^, K^793^, K^796^, K^798^, K^800^, K^803^) mapping within PER2(683-872) was targeted for mutagenesis (Supplementary Figure S4A and B) and recombinant proteins were subjected to *in vitro* ubiquitination in the presence of MDM2 or MDM2(C^470^A) (Figure 3). As shown in Figure 3A, PER2(683-872) is efficiently ubiquitinated by MDM2 (lanes 1-3); however, a form of PER2(683-872) in which all conserved Lys residues were substituted with Ala, named PER2(683-872)-KA (Supplementary Figure S4A), was not targeted for modification (lane 6). This result directly signals that one or more of its Lys residues is a putative target for MDM2-mediated ubiquitination in this fragment (Figure 3A, lanes 3 *vs.* 6).

**Figure 3.**
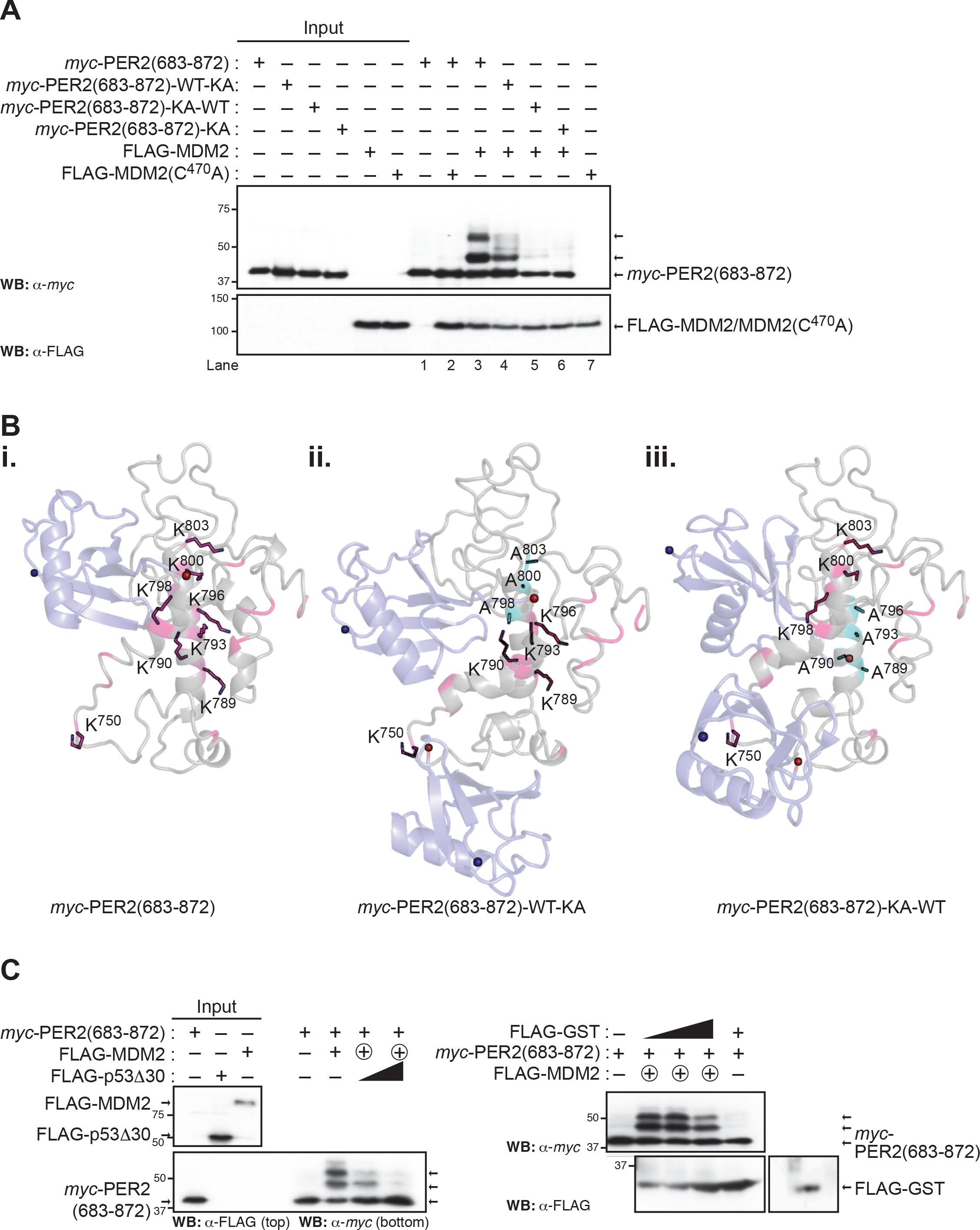
Residues within the central domain of PER2 are ubiquitinated by MDM2. (A) *In vitro* ubiquitination reactions were carried out using recombinant *myc*-tagged PER2(683-872) mutant proteins (WT-KA: K^789,790,793,796^ are wild-type and K^798^A, K^800^A, K^803^A; KA-WT: K^789^A, K^790^A, K^793^A, K^796^A and K^798,800,803^ are wild-type; KA: K^789,790,793,796,798,800,803^A), UbcH5a, and either FLAG-MDM2 or –MDM2(C^470^A). Reaction products were analyzed by immunoblotting using the indicated antibodies. (B) Cartoon representations of PER2(683-872) wild-type, PER2(683-872)-WT-KA, and PER2(683-872)-KA-WT (grey, *panels i, ii, and iii, respectively*). Relevant Lys (K) residues are shown as pink sticks, Lys to Ala mutations are in cyan, and the dominant docked pose of ubiquitin molecules is shown in light blue. In all cases, the C- and N-terminus of ubiquitin are shown in red and blue spheres, respectively. Simulations were performed as indicated in Materials and Methods. (C) Recombinant *myc*-Per2(683-872) and increasing amounts of FLAG-p53Δ30 (comprises residues 1 to 363 of wild-type p53) were pre-incubated and purified before adding (or not, control) FLAG-MDM2. Ubiquitination reactions were allowed to proceed, and samples were analyzed by immunoblotting (*left and right panels*). To rule out non-specific inhibition of Per2(683-872) ubiquitination, FLAG-GST was used as a control (*right panel*). In all cases, molecular weight markers are indicated on the left (kDa), arrows on the right indicate modified forms of the protein substrate, and inputs were 100%. Immunoblot data from **A** and **C** were originated from a single experiment that was repeated three times with similar results.

To gain further insight into the relevance of cluster residues for ubiquitination to take place, two different constructs of PER2(683-872) were generated in which substitutions of K^789^A, K^790^A, K^793^A, and K^796^A were only in PER2(683-872)-KA-WT whereas K^798^A, K^800^A, and K^803^A were solely incorporated into PER2(683-872)-WT-KA (Supplementary Figure S4B). Ubiquitination results showed that, whereas the addition of ubiquitin moieties was reduced in PER2(683-872)-WT-KA, incorporation was completely abrogated in PER2(683-872)-KA-WT (Figure 3A, lanes 3 *vs.* 4 and 5). Although this result might imply that ubiquitination events occur primarily within the upstream mutated Lys residues in PER2(683-872)-KA-WT, we cannot rule out other scenarios including orderly addition of ubiquitin moieties and structural rearrangements, phenotypes that might be disrupted as a result of a single mutation.

To provide insights into possible scenarios, we generated and validated molecular models of PER2(683-872) wild-type, PER2(683-872)-WT-KA, and PER2(683-872)-KA-WT, and carried out unbiased protein-protein docking simulations to predict binding interfaces for ubiquitin molecules in each PER2 fragment (Figure 3B). Unbiased protein-protein docking results strongly favored the placement of a ubiquitin molecule within the K^798^-K^803^ domain with the C-terminus ubiquitin-end making direct contact with K^800^ (Figure 3B, panel *i*). Indeed, modeling and docking predictions in PER2(683-872)-WT-KA suggest that a conformational change would occur and, thus, none of the remaining Lys residues in the fragment, except for K^750^ (outside the two clusters), would be accessible for ubiquitination (Figure 3B, panel *ii*). These findings arise from the analysis of major cluster hits for each protein-ubiquitin docking simulation, the identification of the two most dominant ubiquitin interface poses (both ubiquitin poses are shown in dark blue in each model, Figure 3B, panels *ii* and *iii*), and the comparison of conformational states that show differences between PER2(683-872) and the -WT-KA and/or -KA-WT interfaces.

Modeling results also predicted that it would be unlikely for PER2(683-872)-KA-WT to be ubiquitinated in any Lys residue as a dramatic conformational change precludes the access of ubiquitin to a reactive Lys residue (Figure 3B, panel *iii*). We carried out alanine (Ala) scanning at selected positions by site-directed mutagenesis and determined the contribution of specific Lys residues to PER2 ubiquitination (Supplementary Figure S4C). In agreement with the predicted models and protein-ubiquitin docking results, overall levels of ubiquitination were reduced when single residues, instead of clusters, were replaced by Ala. This was shown more conspicuously for the cases of K^789,790,796,800^ (Supplementary Figure S4C). Further support of our molecular model resulted from ubiquitination experiments carried out using constructs of PER2(683-872) wild type and -WT-KA, where each had the K^750^A mutation (Supplementary Figure S4D). Whereas the former showed a reduction in its ubiquitination status compared to wild-type PER2(683-872), post-translational modification was, as predicted, completely abrogated in the latter (Supplementary Figure S4D).

We previously showed that residues 683 to 872 in PER2 are involved in binding of p53 (22, 24) and, now, that a cluster of Lys residues within that region is targeted for MDM2-mediated ubiquitination (Figure 2B). Consequently, we next asked whether binding of p53 to PER2 could prevent PER2 from being ubiquitinated by MDM2 at the interface of PER2 association with p53. To address this question, we carried out a two-step ubiquitination reaction in which PER2(683-872) was initially incubated with a shorter recombinant version of p53, named p53Δ30 (Figure 3C). We chose to work with p53Δ30 because it lacks the 30 C-terminal residues targeted for ubiquitination by MDM2 but still binds PER2 (22, 43)]. Data showed that addition of increasing amounts of p53Δ30, but not an unrelated protein (GST), to PER2(683-872) gradually decreased MDM2-mediated ubiquitination of the PER2 fragment in the pre-bound complex (Figure 3C). These results further expand the original model in which formation of MDM2:PER2:p53 was proposed to favor p53 stability (24) to include a role for p53 itself in maintaining the integrity of the trimeric complex by preventing PER2 ubiquitination at the interface of their binding.

### MDM2-mediated binding and ubiquitination of PER2 is independent of substrate phosphorylation

Phosphorylation of mouse PER2 on Ser^478^ (Ser^480^ in human PER2) by CK1ε/δ is a prerequisite for β-TrCP binding and subsequent ubiquitination (15, 18); whereas, and despite of its relevance in PER2 stability, priming phosphorylation in Ser^659^ (Ser^662^ in human PER2) and downstream sites are not directly involved in β-TrCP-mediated degradation (9,14,44). Therefore, we asked whether phosphorylation in PER2 is required for MDM2 binding and/or for MDM2 to exert its ubiquitination activity.

Initial binding experiments were performed *in vitro* using recombinantly-tagged proteins in the presence of λPPase, a promiscuous phosphatase enzyme with activity towards phosphorylated serine, threonine, and tyrosine residues in proteins. As shown in Figure 4A, PER2 treatment with λPPase did not abrogate binding to either MDM2 or MDM2(C^470^A); thus, neither phosphorylation in PER2 nor MDM2 E3 ligase activity are a prerequisite for the PER2:MDM2 complex to form. We then specifically ruled out the contribution of CK1ε/δ for PER2 and MDM2 binding by immunoprecipitating the endogenous PER2:MDM2 complex from HCT116 p53^+/+^ cells treated with PF-670462 [named PF670 hereafter, (45)], a specific CK1ε/δ inhibitor with proven effect on circadian rhythms (9, 46) (Figure 4B). We asked whether modification in the critical PER2 Ser^662^ priming site plays a role in PER2-binding to MDM2 and β-TrCP1 ligases. As shown in Supplementary Figure S5A, MDM2 binding to PER2 is independent of priming modifications in Ser^662^ as both PER2 forms, the wild-type and S^662^A mutant, bound MDM2 to the same extent in co-transfected H1299 cells (Supplementary Figure S5A). Further support resulted from immunoprecipitation experiments in which *in vitro* transcribed and translated proteins were allowed to form complexes in HeLa cell extracts before immunoprecipitation. As expected, the S^662^A mutation did not compromise β-TrCP1 binding to PER2 as this site is only relevant to the priming kinase site (Supplementary Figure S5B).

**Figure 4.**
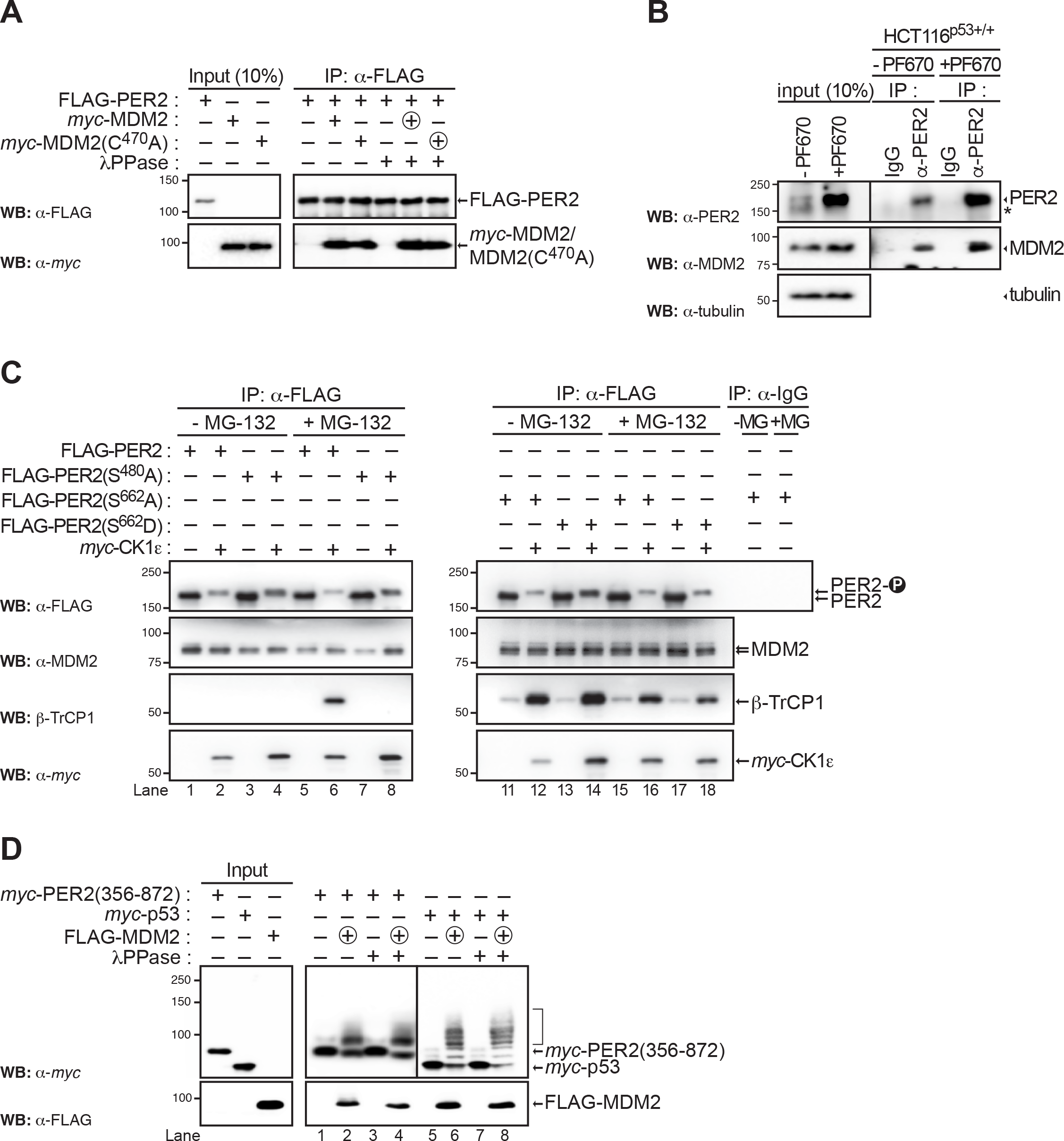
Binding and ubiquitination of PER2 by MDM2 is independent of phosphorylation. (A) *In vitro* transcribed and translated FLAG-PER2 was treated with λPPase and dephosphorylated PER2 was then incubated with either *myc*-MDM2 or *myc*-MDM2(C^470^A) in the presence of binding buffer as indicated in Materials and Methods. Complexes were purified by affinity chromatography and proteins visualized by immunoblotting. (B) Extracts (1 mg) from HCT116 p53^+/+^ cells treated (or not, control) with PF670 inhibitor (1 μM) overnight were immunoprecipitated with α-PER2 antibody (or IgG, control) and protein A-beads. Complexes were resolved by SDS-PAGE and immunoblotting as indicated. Inputs correspond to aliquots (∼100 μg) of total extracts. Asterisk indicates a non-specific band. (C) Cells, HCT116p53+/+, were co-transfected with either empty vector (pCS2+ control, -) or pCS2+^myc^-CK1ε and pCS2+3xFLAG-PER2, -PER2(S^480^A), -PER2(S^662^A), or -PER2(S^662^D). Cells were treated with MG-132 (50 μM, +MG-132 and +MG) or vehicle (control, - MG-132 and -MG) for 4 h before harvesting. Protein complexes were immunoprecipitated using α-FLAG antibody and protein A beads (50% slurry) and samples blotted for endogenous proteins using the indicated antibodies. IgG was used as negative control. (D) *In vitro* transcribed and translated *myc*-PER2(356-872) and p53 were treated with λPPase before carrying out the ubiquitination reaction in the presence or absence (control) of FLAG-MDM2. Reaction products were analyzed by immunoblotting using α-tag antibodies. FLAG-p53 was used as positive control. In all cases, circled “+” symbol indicates proteins added at last. Molecular weight markers are indicated on the left (kDa), arrows on the right indicate modified forms of the protein substrate, and inputs were 100% unless otherwise indicated. Immunoblot data were originated from a single experiment that was repeated twice with similar results.

Next, we asked whether the phosphorylation interplay between the Ser^480^ and Ser^662^ sites would impact the distribution of MDM2 bound to PER2 as previously reported to be the case for β-TrCP1 (9). To explore this scenario, HCT116^p53+/+^ cells were co-transfected with *myc*-CK1ε and FLAG-PER2, FLAG-PER2(S^480^A), PER2(S^662^A), or PER2(S^662^D, mimics phosphorylation in S^662^) constructs and proteins were allowed to express in the presence or absence of the proteasome inhibitor MG132 (Figure 4C). In all cases, FLAG-tagged proteins were immunoprecipitated and endogenous MDM2 and β-TrCP1 bound proteins were detected by immunoblotting. As expected, lower migrating forms of PER2 were detected when co-expressed with CK1ε, a characteristic feature of phosphorylated proteins. In agreement with Figure 4A and B, endogenous MDM2, but not endogenous β-TrCP1, were identified bound to PER2 regardless of CK1ε expression (Figure 4C, lanes 1, 2, 5, and 6). As expected, β-TrCP1 was only detected bound to PER2 when CK1ε was expressed and the proteasome inhibited (Figure 4C, lanes 2 *vs.* 6), whereas its binding was compromised when the Ser^480^ motif in PER2 was altered (Figure 4C, lanes 6 *vs.* 8). Of note, despite the fact that MDM2 and β-TrCP1 were both detected associated to FLAG-PER2 in this assay (Figure 4C, lane 6), our result does not necessarily imply the existence of a trimeric complex that includes each E3 ligase as they can independently associate to PER2 and be simultaneously immunoprecipitated. Our data show MDM2 bound PER2(S^662^A) and PER2(S^662^D) to the same extent confirming that PER2 priming is negligible for MDM2 recognition (Figure 4C, lanes 13-16). As is the case for β-TrCP1, mutation in Ser^662^ favors a larger accumulation of MDM2 protein associated to PER2 (*e.g.*, Figure 4C, lanes 5-6 *vs.* 13-14); however, unlike in the case of β-TrCP in which a larger recruitment of this ligase is associated to a phosphoswitch in PER2 (9), we speculate that greater MDM2 association might result from a structural rearrangement in both forms of mutant PER2. In summary, our results support a model in which recognition of PER2 by MDM2 is independent of phosphorylation in general and in particular by CK1ε and that neither the Ser^480^ nor Ser^662^ phospho-cluster plays a direct role in their recognition.

We also tested whether CK1ε-mediated phosphorylation was, instead, a pre-requisite for PER2 ubiquitination to occur (Supplementary Figure S5C). To evaluate this possibility, we performed a two-step *in vitro* ubiquitination reaction in which the substrate, *myc*-PER2(683-872) was first incubated with FLAG-CK1ε to allow for phosphorylation to occur and then the modified substrate was purified by affinity chromatography before the ubiquitination reaction was carried out in the presence of FLAG-MDM2 (Supplementary Figure S5C, lanes 2-5). Recombinant p53 was used as a control in this experiment (Supplementary Figure S5C, lanes 10-13). As shown, ubiquitination of *myc*-PER2(683-872) was neither abrogated by CK1ε-mediated phosphorylation nor CK1ε binding (Supplementary Figure S5C, lanes 2-9). Pre-treatment of CK1ε with PF670 inhibited the kinase’s activity but not its binding capacity to *myc*-PER2(683-872), and yet, ubiquitination still occurred (Supplementary Figure S5C, lanes 6-9).

To rule out the contribution of phosphorylation events other than those mediated by CK1ε for ubiquitination, *in vitro* reactions were carried out following treatment of various recombinant PER2 fragment proteins with λPPase. As shown for the case of PER2(356-872) and PER2(683-872), and in both Figure 4D and Supplementary Figure S5D, none of the treatments caused a discernible effect in the ubiquitination activity of MDM2 towards its substrate. Lastly, we confirmed binding of CK1ε to *myc*-PER2(356-872) did not sterically block MDM2-mediated ubiquitination of the PER2 fragment (Supplementary Figure S5E). Thus, MDM2-mediated ubiquitination and PER2 phosphorylation seem to follow parallel post-translational paths during PER2’s accumulation in the nucleus.

### MDM2 directly modulates PER2 turnover in cells

As polyubiquitination in proteins is likely a signal for proteasome degradation, we then asked whether MDM2-mediated activity towards PER2 impacts PER2’s half-life. Cells, HCT116^p53+/+^, were transfected with FLAG-MDM2 and harvested at different times after being treated with cycloheximide (CHX), an inhibitor of protein biosynthesis previously used to estimate the half-life of other core clock proteins (23, 47). Analysis of cell lysates showed that endogenous levels of PER2 were remarkably reduced shortly after CHX addition in samples overexpressing MDM2 (Figure 5A, *upper panel*), shortening PER2’s half-life ∼2-fold (Figure 5A, *right graph*). We speculated that a decrease in endogenous MDM2 levels would favor PER2 stability and prolong its half-life, mirroring the effect of β-TrCP1 downregulation on PER2 levels. To test this hypothesis, we transfected HCT116^p53+/+^ cells, which express both MDM2 and β-TrCP1, with *siRNA* targeting either E3 ligase. Lysates were collected at various times after CHX addition and examined for the expression of endogenous levels of PER2, MDM2, and β-TrCP1 by immunoblotting (Figure 5B). Results unmasked two distinct, yet related, features associated with PER2 levels as *siRNA* treatment seemed to influence both PER2’s accumulation and stability (Figure 5B, *upper two panels represent different exposures of the same blot*). First, overall endogenous levels of PER2 increased, albeit at different levels, as a result of knocking down either E3 ligase before CHX addition (Figure 5B, lanes 1 *vs.* 7 and 13). Quantitative analysis showed an ∼2-fold increase in *siMDM2 vs. siβ-TrCP1* treatments and an ∼ 3-fold increase difference compared to mock samples (Figure 5B, lanes 1, 7, and 13 and Supplementary Figure S6). Second, a qualitative assessment of PER2 levels under similar scenarios showed that depletion of either MDM2 or β-TrCP1 stabilized PER2 largely to the same extent (Figure 5B, lanes 7-12 *vs.* 13-18). Overall, our results emphasize the existence of an alternative mode of regulation of PER2 stability.

**Figure 5.**
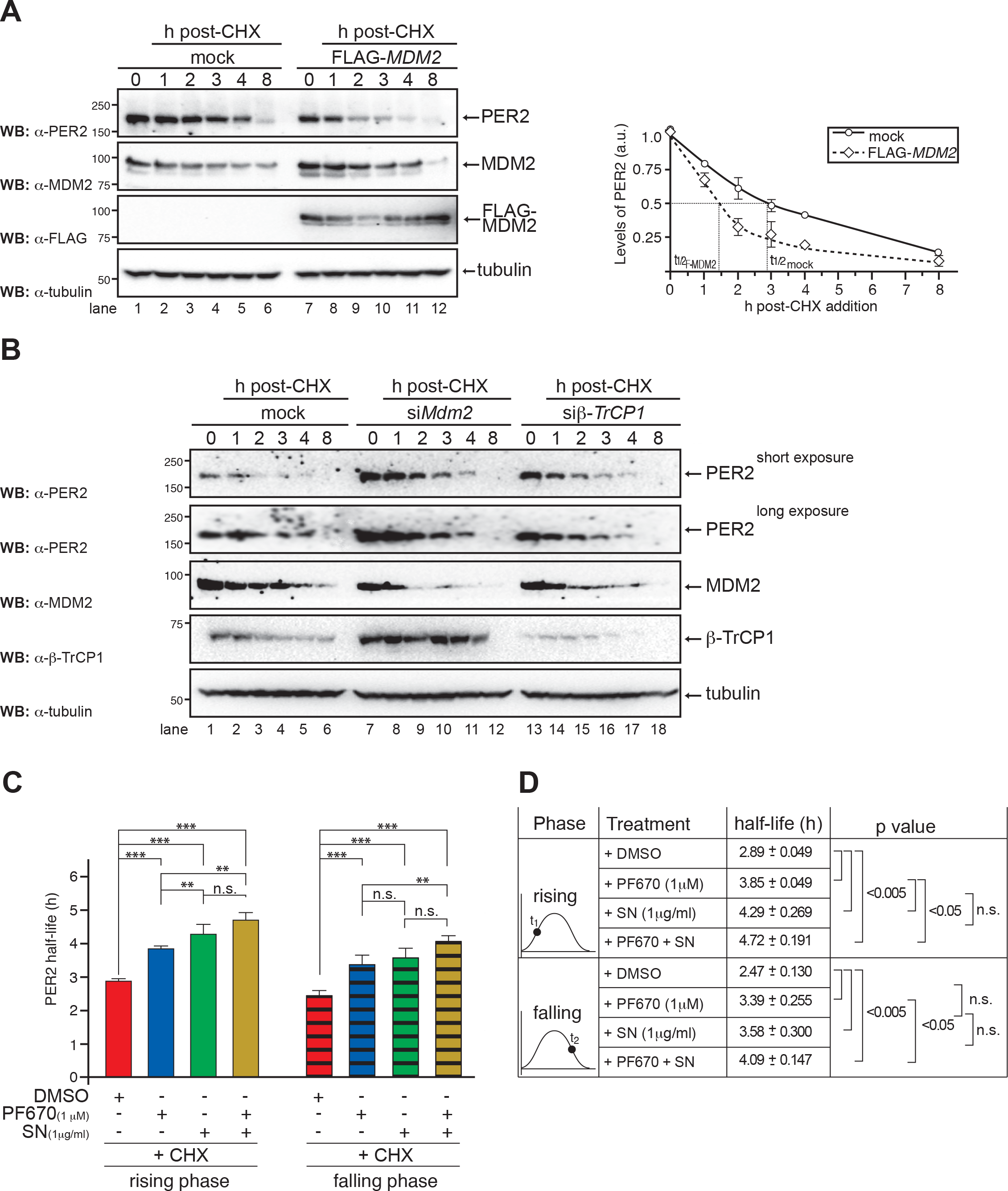
MDM2 modulates PER2 turnover. (**A**) HCT116 cells were transfected with either empty vector (mock) or FLAG-*MDM2* and proteins were allowed to express for 24h before adding cycloheximide (CHX, 100 μg/ml, t=0h). Cells were harvested at different times after CHX addition and extracts were analyzed for endogenous PER2 and MDM2, and FLAG-MDM2 levels by immunoblotting using specific antibodies as indicated on the left. Tubulin was a loading control (*lower panel*). Protein levels of PER2 were quantified using ImageJ Software v1.45 and values normalized to tubulin levels (*right panel*). Immunostaining intensity was plotted as the mean ± SD from three independent experiments. The curve was fitted using Microsoft Excel. (**B**) HCT116 cells were transfected with si-*MDM2* (25 nM), si-*β-TrCP1* (25 nM), or a scrambled siRNA sequence (mock) for 48 h before CHX addition (t=0) as described in M&aterials and Methods. Cells were harvested at different times after CHX addition and extracts were analyzed for the expression of endogenous PER2 (short and long exposures are shown in the *upper two panels*), MDM2, and β-TrCP1 by immunoblotting. Tubulin was a loading control (*lower panel*). (**C**) Real-time bioluminescence recording were carried out in circadian synchronized MEF^mPer2::LUC^ cells maintained for 24h (t1) or 33h (t2) in luciferin-containing media before adding together CHX (40 μg/ml) and DMSO, sempervirine (SN, 1μg/ml), PF670462 (PF670, 1 μM), or a combination of inhibitors. Treatments were performed in triplicate and recordings continue for ∼30h after the addition of inhibitors. Data were normalized to PER2::LUC abundance immediately prior to drug addition. Mean PER2 half-life is shown ± SD. Significance levels were determined by Student’s t test between two groups (*** when *p*<l0.005, ** when *p*<0.05, n.s. means no significant). (**D**) Summary of PER2 protein half-life values obtained under various conditions as described in (**C**) and their statistical significance as determined by t-test.

### Regulation of PER2 stability by MDM2 occurs along the circadian cycle

Next, we asked whether the activity of MDM2 is relevant to PER2 stability during each rising and falling phase of the circadian cycle. Here, we measured real-time bioluminescent rhythms in mouse embryonic fibroblast (MEF) cells in which the mouse *PER2* gene (*mPer2*) was *knocked-in* and the luciferase gene inserted downstream [named MEF^mPer2::LUC^ hereafter, (48)]. Studies confirmed that MEF^mPer2::LUC^ cells maintained robust rhythms in luciferase activity for several days and the mPer2::LUC fusion protein showed rhythms of accumulation and posttranslational modifications that mirrored those described *in vivo* (48, 49).

First, degradation of mPER2::LUC was monitored by bioluminescence recordings in cells treated with CHX and sempervirine nitrate, a compound that specifically inhibits the ubiquitin ligase activity of MDM2 [named SN hereafter, (50, 51)] at either rising (t_1_) or falling phases (t_2_) (Figures 5C and D). Inhibition of MDM2 activity resulted in a longer PER2 half-life compared to vehicle (DMSO) suggesting a role for the E3 ligase in modulating PER2 stability in both circadian phases [Figures 5C and D; t_1_(DMSO *vs.* SN, in h): 2.89 ± 0.049 *vs.* 4.29 ± 0.269 *p*<0.005 and t_2_(DMSO *vs.* SN, in h): 2.47 ± 0.130 *vs.* 3.58 ± 0.300 *p*<0.005]. As expected, an increase in PER2’s half-life was noticeable when cells were treated with PF670 as CK1δ/ε-mediated phosphorylation of Ser^480^ is precluded and, therefore, β-TrCP1 was unable to recognize its substrate [Figures 5C and D; t_1_(DMSO *vs.* PF670, in h): 2.89 ± 0.049 *vs.* 3.85 ± 0.049, *p*<0.005 and t_2_(DMSO *vs.* PF670, in h): 2.47 ± 0.130 *vs.* 3.39 ± 0.255, *p*<0.005 and (15)]. These results establish the relevance of MDM2 activity for modulating PER2 stability during the accumulation and degradation phases of the circadian cycle.

We then evaluated how SN and PF670 treatments compared to each other in terms of mPER2::LUC stability in both rising and falling phases (Figure 5C). Results showed a marginal, but consistent, significant increase in mPER2::LUC half-life between SN and PF670 treatments only when administered during the rising phase [Figures 5C and D; t_1_(PF670 *vs.* SN, in h): 3.85 ± 0.049 *vs.* 4.29 ± 0.269, *p*<0.05 and t_2_(PF670 *vs.* SN, in h): 3.39 ± 0.255 *vs.* 3.58 ± 0.300, n.s.]. In this context, our results suggest that MDM2 and β-TrCP are both needed during the circadian accumulation phase of PER2 but might have redundant roles during the falling phase where no significant difference in PER2 half-life was observed between treatments. Consequently, it seemed relevant to explore the contribution of CK1ε/δ for PER2 accumulation in the context of MDM2 activity.

Cells, MEF^mPer2::LUC^, were then incubated with a combination of SN and PF670 inhibitors in the presence of CHX as described in the Methods section. Remarkably, the addition of both inhibitors had a synergistic effect on the stability of mPER2::LUC when compared to the sole addition of PF670, but not SN, in either phase [Figure 5C; t_1_(PF670 *vs.* PF670+SN, in h): 3.85 ± 0.049 *vs.* 4.72 ± 0.191, *p*<0.05 and t_2_(PF670 *vs.* PF670+SN, in h): 3.39 ± 0.255 *vs.* 4.09 ± 0.147, *p*<0.05]. Overall, our data supports a model where PER2 stability depends, *a priori*, on the interplay between both E3 ligases.

### MDM2’s function is required to maintain circadian period

Maintenance of circadian oscillations relies on the expression of the rate-limiting component PER2 for the formation of a functional PER2:CRY inhibitory complex (49). Therefore, we hypothesized that alterations in PER2 stability that result from tuning MDM2’s levels or activity should impact the length of the circadian period.

Our initial studies focused on measuring the period length of the circadian oscillator in MEF^mPer2::LUC^ cells in which MDM2’s level was either augmented by its overexpression or silenced by *siRNA* targeting (Figures 6A and B and Supplementary Figures S7A and C). As expected from our biochemical findings (Figure 5), synchronized MEF^mPer2::LUC^ cells overexpressing MDM2 exhibited a shorter period length (25.20 ± 0.100 h *vs.* 24.53 ± 0.05 h, *p*<0.005) compared to mock-transfected cells (Figure 6A and Supplementary Figure 7E). Furthermore, transfections of MEF^mPer2::LUC^ with increasing amounts of *myc*-MDM2 significantly resulted in a dose-dependent shortening of circadian period length by up to ∼1.5h even at low levels of *myc*-MDM2 transfection (Supplementary Figure 7B), suggesting that a tight regulation of MDM2 needs to be maintained under physiological conditions to ensure proper oscillation. Next, we challenged the model by hypothesizing that knockdown expression of MDM2 by siRNA transfection (named siMDM2) of MEF^mPer2::LUC^ cells should result in the converse phenotype and, thus, a lengthened period (Figure 6B and Supplementary Figure 7C). Indeed, our data showed that downregulation of MDM2 expression resulted in significant lengthening of the circadian period (25.50 ± 0.141 h *vs.* 26.75 ± 0.480 h, *p*<0.05, Supplementary Figure 6E), confirming the requirement of MDM2 for normal circadian oscillations.

**Figure 6.**
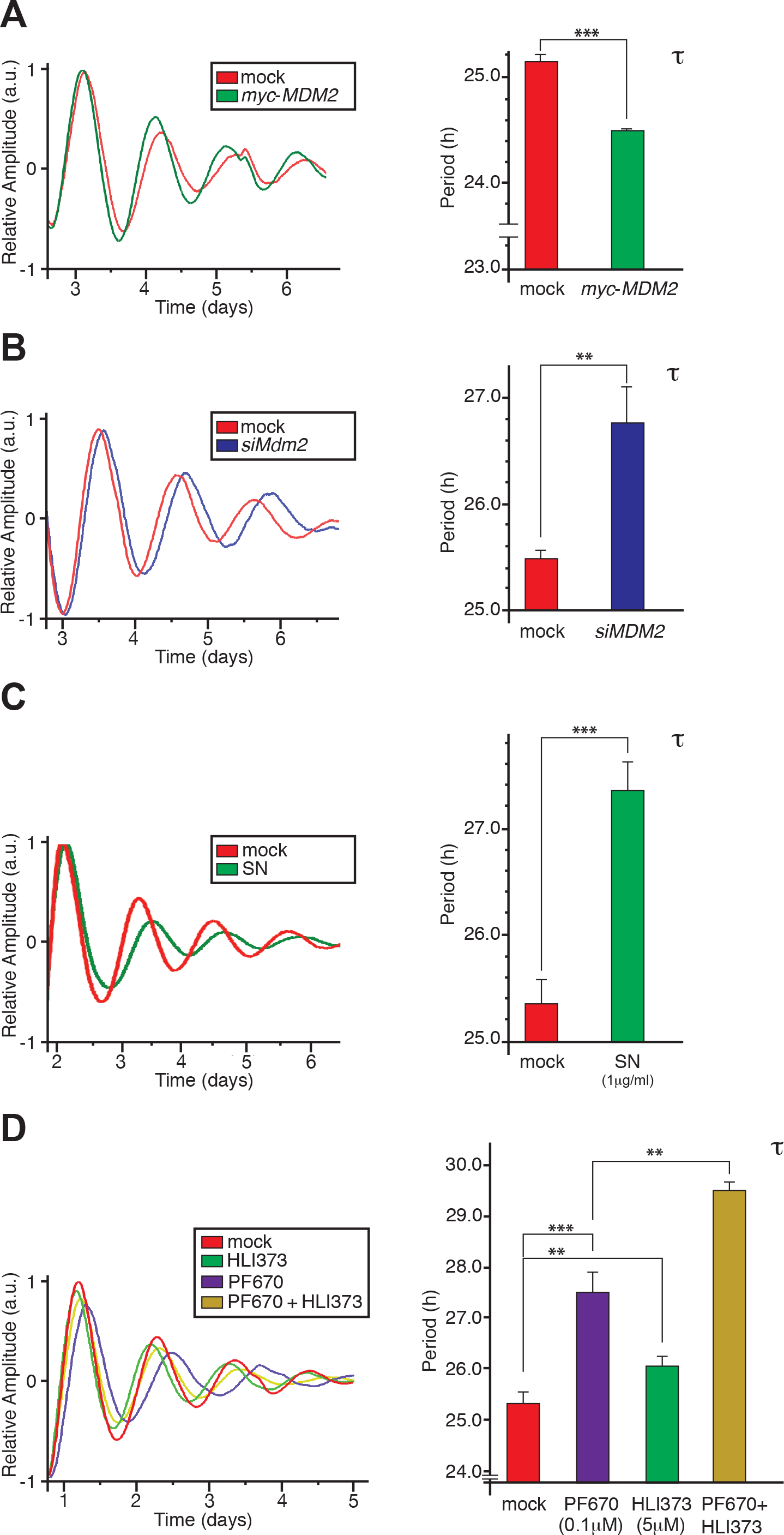
The activity of MDM2 influences the length of the circadian period. (**A**) MEF^mPer2::LUC^ cells were transfected with either pCS2+-*myc* (mock) or pCS2+-*myc*-MDM2 and proteins were allowed to express for 24h before cells were circadian synchronized. Abundance of PER2::LUC was monitored by bioluminescence for 7 days as described in Materials and Methods (*left panel*). In other experiments, MEF^mPer2::LUC^ cells were transfected with *siMDM2* [(**B**), a scramble siRNA was used as mock control] or treated with sempervirine [SN, 1μg/ml, (**C**)] before synchronization and maintained after. (**D**) Synchronized MEF^mPer2::LUC^ cells were incubated with HLI373 (5μM), PF670462 (PF670, 0.1 μM), or acombination of both inhibitors. Cells were maintained with inhibitors at all times during data collection. Biological replicates were carried out in triplicate. For **A-D**, bar graphs indicate the length of the circadian period calculated using the LumiCycle analysis software (Actimetrics). The vehicle, DMSO, was used as control (mock). Values are the mean ± SD from three independent experiments. Statistical significance was determined by t-test. \*\*\**p* ≤ 0.005, \*\**p*≤ 0.05.

Following, MEF^mPer2::LUC^ cells were treated with the cell permeable MDM2 inhibitor SN or HLI373 at a dose that *i*) prevented MDM2 auto-ubiquitination and degradation (51–53), and *ii*) did not affect cell viability (Supplementary Figures 7D and E). Synchronized MEF^mPer2::LUC^ cells were maintained in the presence of the inhibitor throughout the time course during bioluminescence recording (Figure 6C and Supplementary Figures S7G). Average bioluminescence rhythms of SN-treated cells show a dramatic lengthening of the circadian period of ∼2 h (25.35 ± 0.311 h *vs.* 27.33 ± 0.340 h, *p*<0.0005, Supplementary Figures 7G), which closely resembled the result obtained when transfecting MEF^mPer2::LUC^ cells with siMDM2 (Figures 6B *vs.* C), suggesting that control over MDM2’s activity remains a major point of regulation. Similarly, a delay in period length was also observed when MEF^mPer2::LUC^ cells were treated with the HLI373 inhibitor (mock: 24.97 ± 0.208 h, HLI373: 26.10 ± 0.283, h *p*<0.05, Supplementary Figure S7F). Therefore, despite the fact that changes in MDM2 levels influence circadian oscillations, MDM2’s E3 ligase activity is actually the chief contributor to the observed phenotype.

Then, we evaluated whether the combined effect of PF670 and HLI373 on PER2 stability results in a synergistic change in circadian lengthening (Figure 6D). In this scenario, synchronized MEF^mPer2::LUC^ cells were maintained with either inhibitor (PF670 or HLI373) or a combination of both (PF670+ HLI373) and the long-term effect in bioluminescence rhythms were simultaneously recorded throughout the time course analyzed. In agreement with Figure 5C, and (9) for PF670, treatment of MEF^mPer2::LUC^ cells with HLI373 resulted in a significant increase in circadian period length even a low concentrations (mock: 25.35 ± 0.071 h; HLI373: 26.10 ± 0.200 h; PF670: 27.53 ± 0.404 h, PF670+HLI373: 29.50 ± 0.173 h, Supplementary Figure S7G). Similarly, treatment of MEF^mPer2::LUC^ cells with SN alone or in combination with PF670, also exhibited period lengthening, further advocating for the specific involvement of MDM2’s activity in circadian oscillation. Our findings showed that, although significant, the effect of both inhibitors is not additive but synergistic and reflects on the cell’s period length. Overall, our data suggest a model in which both events, ubiquitination of PER2 by MDM2 and PER2 phosphorylation by CK1ε/δ, are relevant to determining the circadian period length despite the appearance that both events seem, *a priori*, to take place independently of each other.

## DISCUSSION

Timely degradation of regulatory proteins is essential for most aspects of cellular homeostasis and is relevant to signaling processes involved in cell growth, proliferation, and survival. It is, therefore, not surprising that malfunctioning of any aspect of the protein degradation process results in a wide spectrum of diseases and disorders [for review see (54)]. Because mammalian circadian rhythm also relies on the continuous cycle of protein synthesis and degradation, it is not exempted from problems associated with protein turnover dysregulation. Indeed, mice bearing loss-of-function mutations or knock-down expression in genes encoding ubiquitin-modifying enzymes involved in clock regulation (*e.g.*, *FBXL3*, *FBXL21*, *FBW1A*, *HUWE1*, *PAM*, *UBE3A*, *SIAH2*) exhibit a phenotype where the free-running period of locomotor activity is longer, shorter, or dampened [for review see (55)]. As clock components control the expression of an array of genes involved in multiple cellular mechanisms, it is reasonable to expect that alteration in their expression and accumulation is linked to various human diseases (56). This brings into consideration the relevance of PER2, a circadian component whose function lies at the intersection of the cellular response to DNA-damage (24) and whose turnover depends on its phosphorylation by CK1ε/δ and β-TrCP1/2 binding, followed by ubiquitination and proteasomal degradation [see (8) and references within]. Whereas substantive research has made a compelling case for how PER2 accumulates and how its level modulates the function of the clock, it poses the question of whether PER2 turnover remains exclusively a β-TrCP1/2 matter. And, whereas the answer could have certainly been affirmative, seemingly unnoticed observations suggested to us that alternative scenarios could exist. For example, the counterintuitive finding that PER2’s half-life was reported to be shorter in cultured cells co-expressing the dominant negative forms of β-TrCP1 and 2 (β-TrCP1ΔF and 2 ΔF) (57), the need for phosphorylation-independent mechanisms of PER2 degradation to exist to explain its three-stage kinetics of degradation (9), or the presence of ubiquitinated forms of PER2 in a biological system in which β-TrCP1 and 2 are knocked down (21). As a result, we turned to our findings that established that PER2 is able to form a trimeric complex with MDM2 and p53 (22) and asked whether MDM2 might play a role in PER2 stability.

Our results show that PER2 binds to MDM2 (PER2:MDM2) in a p53-independent manner *in vitro* and exists as a readily detectable endogenous complex in various cell settings (Figure 1 and Supplementary Figure 1). This is a non-trivial finding as, to the best our knowledge, all well-established E3 ubiquitin ligases acting on clock components only recognize phosphorylated substrates. This includes, in addition to β-TrCPs, the E3 ligases FBXL3 and FBXL21, which act on AMPK-mediated phosphorylated CRY1/2 (58), FBXW7, which acts on cyclin-dependent kinase 1-mediated phosphorylated REV-ERBα (59), and a yet uncharacterized E3 ligase that targets GSK3β-mediated phosphorylated BMAL1 (60). The challenge of identifying novel E3 ubiquitin ligases targeting clock components has led to the development of screenings that revolve around identifying enzyme-substrate binding or functional interactions (61, 62). Of these, the recent identification of the ubiquitin ligase Siah2, which regulates REV-ERVα turnover, has been the most promising finding, yet, it belongs to the domain of ligases that recognize phosphorylated substrates (62).

Binding of PER2 to MDM2 occurs in a region distinct from those identified for p53 binding and E3 ligase activity (Figures 1-2), a result in agreement with the existence of the PER2:MDM2:p53 complex (22). Next, we established PER2 as a novel substrate of MDM2 and, conversely, MDM2 resulted in a previously uncharacterized E3 ligase responsible for PER2 ubiquitination (Figure 2). The relevance of these initial findings lays in the existence of an alternative mechanism to recognize and target PER2 for degradation that is independent of phosphorylation (Figures 2-4).

Binding specificity among E2 enzymes of the UbcH5 family by MDM2 defined the intrinsic preference for K^11^, K^48^, and K^63^ ubiquitin linkages in PER2 resulting in the incorporation of multi-ubiquitin molecules (Figures 2-3). Docking of K^11,48, and 63^ ubiquitin conformations favored elongation and, thus, the formation of poly-ubiquitinated chains (42). Although the addition of multiple ubiquitination units in PER2 by MDM2 represents a novel finding among circadian proteins, post-translational modifications of this nature by MDM2 are not uncommon as shown, for example, in the case of p53 and FOXO4 (40, 63).

We found that the accumulation and half-life of endogenous PER2 varied in scenarios in which MDM2 levels or activity were modulated, which also altered the cell’s circadian period length (Figures 5-6). These findings raise the question of how MDM2’s primary role as a PER2 regulator would fit into the functioning of the actual mammalian clock mechanism when acting under normal physiological conditions. This is certainly a difficult question to address, especially considering that MDM2 distribution in normal cells is largely nuclear, that MDM2 could promote either mono-or poly-ubiquitination of substrates depending on its endogenous levels, and that rhythmic levels of MDM2 protein and transcripts are largely absent in unstressed cells [for review see (64) and (23)]. Whereas these well-established premises create constraints around the possible function of MDM2 within the clock molecular mechanism, we propose a few scenarios for further consideration. For example, it is possible that, under physiological conditions, translocation of PER2 to the nucleus would initially result in time-of-day accumulation of CK1ε/δ-dependent phosphorylation events in PER2 that may serve to prime the substrate first for β-TrCP1/2-mediated degradation and later for MDM2 targeting. Furthermore, it is not uncommon to find that the generation of polyubiquitination substrates targeted for proteasomal degradation require both priming of mono-ubiquitinated substrates and intrinsic E3 ligase activity of more than one enzyme as has been shown, for example, in the case of p53 (65, 66).

Phosphorylation of PER2 by CK1ε/δ either stabilizes or destabilizes the circadian factor depending on the phosphocluster targeted in PER2, and thus, adjusts the length of the circadian period to diverse environmental stimuli [*e.g.*, temperature, metabolic signal, (9, 11)]. Our findings open the possibility of PER2’s stability being modulated by signals that converge in MDM2, for example, those that respond to genotoxic and cytotoxic cellular stress, and for which a change in period length might provide a fitness advantage. Indeed, MDM2’s activity can be modulated by post-translational modifications, stability, localization, or binding and be exquisitely tuned by, for example, alteration in oxygen levels, exposure to low-dose radiation, and even slight changes in growth factor concentrations (64). Certainly, phase resetting of the mammalian circadian clock has been shown to occur in response to DNA-damage and metabolic stress in both cell cultures and animal models, a phenotype that is increasingly associated with the existence of crosstalk mechanisms between clock proteins and checkpoint components (24,67–69).

At this point, the role of MDM2:PER2 interaction in the mammalian system and within any of the scenarios described above remains largely within the domain of speculation and represents an area of active research in our laboratory. We expect that mounting biochemical, molecular, and genetic evidence will provide a conceptual framework within which we can understand how cells relate and respond to environmental perturbations, no longer in isolation, but in the context of multicellular systems.

## ACKNOWLEDGEMENTS

This article is dedicated by C.V.F. to the memory of Dr. James Maller, Professor and Howard Hughes Investigator, a pioneer in the field of cell cycle regulation and a fine mentor. The authors thank Dr. D. G. S. Capelluto and Dr. J. Tyson for critical reading of the manuscript and all members of the Finkielstein laboratory for help and discussions. We would also like to thank Dr. Shihoko Kojima for reagents and advice and Dr. J. Webster for comments and manuscript editing. J.L. performed all experiments and statistical analyses discussed in this article except those specifically mentioned below. X.Z. carried out the experiments shown in Figures 3C, 4, 5C-D, 6D, S3B, S4C and D, S5C-E, S6D, and S7D. T.G. contributed with Figures 2, 3A and C, S2A and C, S3A and C, S4A. L.J. contributed with Figure S7E. A.B. performed and analyzed the modeling shown in Figure 3B. J.L., X.Z., and J.K.K. analyzed the overall data and contributed to refining the hypothesis. C.V.F. and J.L. conceived this project. C.V.F. supervised and coordinated all investigators for the project and wrote the manuscript.

## FUNDING

This work was supported by the National Science Foundation MCB division (MCB-1517298) and the Fralin Life Science Institute to C.V.F. and the National Research Foundation of Korea (N01160447) to J.K.K.

## SUPPLEMENTARY FIGURE LEGENDS

**Figure S1. *In vitro* binding studies of MDM2.** (**A**) H1299 cells were co-transfected with pCS2+*myc*-PER2, FLAG-MDM2, FLAG-MDM2(C^470^A), or FLAG-β-TrCP1 and extracts were immunoprecipitated and bound proteins analyzed by immunoblotting using specific antibodies. (**B**) Schematic representation of MDM2 constructs [Uniprot ID: Q00987-11, MDM2(1-117), MDM2(1-230), MDM2(1-434), MDM2(117-497), MDM2(230-497), MDM2(434-497)] used in binding mapping experiments. All MDM2 constructs encode an N-terminus FLAG-tag. Structural and functional domains in MDM2 are indicated as boxes in the full-length representation. NES, nuclear export signal; NLS, nuclear localization signal. Epitope mapping of specific MDM2 antibodies are indicated as: 4B2, comprises residues 19-50; SMP14, comprises residues 154-167; and 4B11, comprises residues 383-491. (**C**) *In vitro* transcribed and translated FLAG-MDM2 recombinant proteins were incubated with *myc*-PER2 and the complex was allowed to form before samples were immunoprecipitated using α-FLAG antibody and protein A beads as indicated in Materials and Methods. The complex was then washed with increasing concentration of NaCl (100 mM, 250 mM, and 500 mM) and bound PER2 was detected using α-*myc* antibody. In all cases, molecular weight markers are indicated on the left (kDa). Asterisk indicates non-specific band. Unless indicated, all experiments were repeated at least twice with similar results.

**Figure S2. Identification of ubiquitinated forms of PER2 in cells.** (**A**) *In vitro* ubiquitination reactions were carried out using recombinant tagged full-length PER2 and MDM2/MDM2(C^470^A) proteins in the presence of FLAG-ubiquitin and E1/E2 ubiquitin enzymes. Reactions were allowed to proceed and the modified PER2 substrate was purified by immunoprecipitation. Samples were resolved by SDS-PAGE and immunoblotted using α-FLAG (*upper panel*) or α-*myc* antibodies (*middle* and *lower panels*). (**B**) Cells, HCT116^p53+/+^, were co-transfected with FLAG-PER2, *myc*-MDM2, *myc*-MDM2(C^470^A), or empty vector (control, -) and maintained in the presence (+) or absence (-) of MG132. Cells were harvested and PER2 ubiquitination detected by immunoprecipitation using α-FLAG antibody and immunoblotting with α-ubiquitin (*upper panel*), α-PER2 (*middle panel*), or α-*myc* (*lower panel*) antibodies. (**C**) An *in vitro* transcribed and translated *myc*-tagged fragment of PER2 comprising residues 1-682 was assessed for ubiquitination *in vitro* using either FLAG-MDM2 or -MDM2(C^470^A) ligases. Products were identified by immunoblotting using the indicated antibodies. Inputs were 100%. Bracket indicates ubiquitinated forms of PER2(1-682). In all cases, molecular weight markers are indicated on the left (kDa). Asterisk indicates non-specific band.

**Figure S3. PER2’s ubiquitination is tightly regulated by MDM2.** (**A**) The *in vitro* ubiquitination reaction of *myc*-PER2(683-872) by FLAG-MDM2 was optimized for the levels of its co-factor and co-substrate, ATP and ubiquitin, to maximize the incorporation of ubiquitin moietis. The *myc*-PER2(683-872):FLAG-MDM2 complex was allowed to form before adding the ubiquitin mix containing different 1- to 3-fold level increases of ATP and/or ubiquitin as described in Materials and Methods using UbcH5a as E2 enzyme. Ubiquitinated forms of *myc*-PER2(683-872) were detected using α-*myc* antibody (*upper panel*) and levels of FLAG-MDM2 confirmed by immunoblotting using α-FLAG antibody (*lower panel*). (**B**) A time-course *in vitro* ubiquitination assay was carried out using the condition optimized in (**A**) and either FLAG-MDM2 or FLAG-MDM2(C^470^A) enzymes. Samples were taken before incubation (t=0) and at various times after as indicated. Ubiquitinated forms of the substrate were detected by immunoblotting using α-*myc* antibody (*upper panel*). Levels of each form of E3 ligase were assessed by blotting using α-FLAG (*middle and lower panels*). (**C**) *Left panel.* Recombinant *myc*-PER2(683-872) was subject to ubiquitination in a reaction containing E1, one of the indicated forms of E2 enzyme UbcH5 (a, b, c), and either FLAG-MDM2 or –MDM2(C^470^A) and the products were analyzed by immunoblotting. The FLAG-p53 substrate was included as control (*right panel*). Poly-ubiquitinated forms of p53 are indicated between brackets on the right. Arrows on the right indicate substrate modification, and molecular weight markers are indicated on the left (kDa).

**Figure S4. Identification of putative ubiquitination sites within the PER2 central domain.** (**A**) Sequence alignment of a central region of the human PER2 protein corresponding to residues 789 to 806 (*H. sapiens*, Accession NP_073728) with a comparable region from *M. musculus* (Accession NP_035196), *R. norvegicus* (Accession NP_113866), *B. Taurus* (Accession XP_589710), *G. gallus* (Accession NP_989593), *D. rerio* (Accession NP_878277), and *X. laevis* (Accession NP_001081098). (**B**) Table summarizing all seven conserved Lys residues (K^789^, K^790^, K^793^, K^796^, K^798^, K^800^, K^803^) mutated to Ala (indicated as A) in the four different constructs (WT, WT-KA, KA-WT, KA) used for the assay in Figure 3A. (-) indicates Lys remains as wild-type residue. (**C**) *In vitro* ubiquitination reactions were carried out using recombinant forms of *myc*-PER2(683-872) where single Lys residues were substituted by Ala. Reactions were allowed to proceed in the presence of wild-type MDM2 or its enzymatically inactive form MDM2(C^470^A). (**D**) Reactions were as in (**C**) except two substrates bearing an additional mutation K^750^A, *myc*-PER2(683-872)-K^750^A and *myc*-PER2(683-872)-WT-KA-K^750^A, were included in the assay. Arrows on the right indicate substrate modification, and molecular weight markers are indicated on the left (kDa).

**Figure S5. Ubiquitination of the central domain of PER2 is independent of phosphorylation.** (**A**) H1299 cells were co-transfected with pCS2+*myc*-PER2, *myc-*PER2(S^662^A), pCS2+3xFLAG-MDM2, or 3xFLAG-MDM2(C^470^A). Cell extracts were incubated with α-FLAG antibody and protein A beads (50% slurry) and bound proteins were identified by immunoblotting using α-tag antibodies. (**B**) *In vitro* transcribed and translated tagged proteins (lanes 1 to 4) were mixed and complexes allowed to form before immunoprecipitation. Bound components were identified by immunoblotting using the indicated antibodies. (**C**) Phosphorylation of *myc*-PER2(683-872) was carried out in a two-step process in which the recombinant PER2 protein was pre-incubated with FLAG-CK1ε and the phosphorylated substrate was subjected to ubiquitination using FLAG-MDM2. In other reactions, FLAG-CK1ε was pre-incubated with PF670 before adding the components to the kinase reaction. Products were analyzed by immunoblotting using specific antibodies. DMSO was a vehicle control. Circled “+” symbol indicates proteins added at last and after the immunoprecipitated complex was purified and washed. FLAG-p53 was used as positive control. (**D**) Recombinant *myc*-Per2(683-872) was incubated (or not, control) with lambda phosphatase (λPPase, 400U/μl) to remove any phosphate group incorporated in *myc*-Per2(683-872) before adding FLAG-MDM2 (indicated as a circled “+”). Ubiquitination reactions were carried out and products resolved as indicated in Materials and Methods using α-*myc* (*upper panels*) or α-FLAG antibodies (*lower panel*). (**E**) The *myc*-Per2(356-872) recombinant protein was pre-incubated with increasing amounts of CK1ε and the complexes were allowed to form before the ubiquitination reaction was carried out in the presence of FLAG-MDM2. In all cases, arrows on the right indicate ubiquitinated forms of each fragment and molecular weight markers are indicated on the left (kDa). Unless indicated, inputs were 100%. Immunoblot data were originated from a single experiment that was repeated twice times with similar results.

**Figure S6. Quantification of endogenous protein levels.** Protein levels of PER2, MDM2, and β-TrCP1 were quantified using ImageJ Software v1.45 from the experiment depicted in Figure 5B and values normalized to tubulin levels. Immunostaining intensity was plotted as the mean ± SD from three independent experiments.

**Figure S7. MDM2’s level and activity influence circadian period length.** (**A**) In a parallel set of dishes, pCS2+*myc*-MDM2 transfected MEF^mPer2::LUC^ cells were monitored for either PER2::LUC luminescence activity (Figure 6A) or MDM2 protein expression. MDM2 was detected in lysates (40 μg) 24 h after transfection (t=-2), 2h after dexamethasone treatment (t=0), and at various times after synchronization (t=24, 40, 40.5, and 52 h) by immunoblotting using an α-*myc* antibody. “Mock” indicates cells transfected with empty plasmid. (**B**) MEF^mPer2::LUC^ cells were transfected with various amounts of pCS2+-*myc*-MDM2, synchronized with dexamethasone, and monitored for luminescence activity over time (*left panel*). The circadian period length for each treatment was determined using LumiCycle analysis software (*right panel*). (**C**) Knockdown expression of MDM2 in MEF^mPer2::LUC^ cells was confirmed by immunoblotting of lysates collected 24 and 48 h after siMDM2 transfection (25 nM), following dexamethasone addition (t=0), and at different times after synchronization (t=72 and 96 h). (**D**) Cell viability was assayed in MEF^mPer2::LUC^ cells incubated with different concentrations of sempervirine (SN) for up to 7 days using the MTT cell viability kit and following manufacturer’s instructions (ThermoFisher). Values are the mean ± SD from three independent experiments repeated in triplicate. (**E**) Cell viability test was performed as in (**D**) but using the HLI373 inhibitor (5 μM) instead. Values are the mean ± SD from three independent experiments repeated in triplicate. (**F**) Synchronized MEF^mPer2::LUC^ cells were incubated with HLI373 (5 μM) and maintained with the inhibitor at all times during data collection. The circadian period length was calculated using the LumiCycle analysis software (Actimetrics). DMSO was used as control (mock). (**G**) Summary of circadian period length data obtained from the various treatment modalities included in Figure 6. Values are the mean ± SD from three independent experiments. Statistical significance between two groups was determined by t-test.

## REFERENCES

1. Masri, S., Cervantes, M. and Sassone-Corsi, P. (2013) The circadian clock and cell cycle: interconnected biological circuits. Curr Opin Cell Biol, 25, 730–734.

2. Bell-Pedersen, D., Cassone, V.M., Earnest, D.J., Golden, S.S., Hardin, P.E., Thomas, T.L. and Zoran, M.J. (2005) Circadian rhythms from multiple oscillators: lessons from diverse organisms. Nat Rev Genet, 6, 544–556.

3. Buhr, E.D. and Takahashi, J.S. (2013) Molecular components of the Mammalian circadian clock. Handbook of experimental pharmacology, 3–27.

4. Yagita, K., Yamaguchi, S., Tamanini, F., van Der Horst, G.T., Hoeijmakers, J.H., Yasui, A., Loros, J.J., Dunlap, J.C. and Okamura, H. (2000) Dimerization and nuclear entry of mPER proteins in mammalian cells. Genes Dev, 14, 1353–1363.

5. Ripperger, J.A. and Albrecht, U. (2012) The circadian clock component PERIOD2: from molecular to cerebral functions. Prog Brain Res, 199, 233–245.

6. Lee, H.M., Chen, R., Kim, H., Etchegaray, J.P., Weaver, D.R. and Lee, C. (2011) The period of the circadian oscillator is primarily determined by the balance between casein kinase 1 and protein phosphatase 1. Proceedings of the National Academy of Sciences of the United States of America, 108, 16451–16456.

7. Chiu, J.C., Ko, H.W. and Edery, I. (2011) NEMO/NLK phosphorylates PERIOD to initiate a time-delay phosphorylation circuit that sets circadian clock speed. Cell, 145, 357–370.

8. Gallego, M. and Virshup, D.M. (2007) Post-translational modifications regulate the ticking of the circadian clock. Nat Rev Mol Cell Biol, 8, 139–148.

9. Zhou, M., Kim, J.K., Eng, G.W., Forger, D.B. and Virshup, D.M. (2015) A Period2 Phosphoswitch Regulates and Temperature Compensates Circadian Period. Molecular cell, 60, 77–88.

10. Kaasik, K., Kivimae, S., Allen, J.J., Chalkley, R.J., Huang, Y., Baer, K., Kissel, H., Burlingame, A.L., Shokat, K.M., Ptacek, L.J. et al. (2013) Glucose sensor O-GlcNAcylation coordinates with phosphorylation to regulate circadian clock. Cell Metab, 17, 291–302.

11. D’Alessandro, M., Beesley, S., Kim, J.K., Chen, R., Abich, E., Cheng, W., Yi, P., Takahashi, J.S. and Lee, C. (2015) A tunable artificial circadian clock in clock-defective mice. Nature communications, 6, 8587.

12. Meng, Q.J., Logunova, L., Maywood, E.S., Gallego, M., Lebiecki, J., Brown, T.M., Sladek, M., Semikhodskii, A.S., Glossop, N.R., Piggins, H.D. et al. (2008) Setting clock speed in mammals: the CK1 epsilon tau mutation in mice accelerates circadian pacemakers by selectively destabilizing PERIOD proteins. Neuron, 58, 78–88.

13. Lowrey, P.L., Shimomura, K., Antoch, M.P., Yamazaki, S., Zemenides, P.D., Ralph, M.R., Menaker, M. and Takahashi, J.S. (2000) Positional syntenic cloning and functional characterization of the mammalian circadian mutation tau. Science, 288, 483–492.

14. Shanware, N.P., Hutchinson, J.A., Kim, S.H., Zhan, L., Bowler, M.J. and Tibbetts, R.S. (2011) Casein kinase 1-dependent phosphorylation of familial advanced sleep phase syndrome-associated residues controls PERIOD 2 stability. The Journal of biological chemistry, 286, 12766–12774.

15. Eide, E.J., Woolf, M.F., Kang, H., Woolf, P., Hurst, W., Camacho, F., Vielhaber, E.L., Giovanni, A. and Virshup, D.M. (2005) Control of mammalian circadian rhythm by CKIepsilon-regulated proteasome-mediated PER2 degradation. Mol Cell Biol, 25, 2795–2807.

16. Ohsaki, K., Oishi, K., Kozono, Y., Nakayama, K., Nakayama, K.I. and Ishida, N. (2008) The role of {beta}-TrCP1 and {beta}-TrCP2 in circadian rhythm generation by mediating degradation of clock protein PER2. J Biochem, 144, 609–618.

17. Fuchs, S.Y., Spiegelman, V.S. and Kumar, K.G. (2004) The many faces of beta-TrCP E3 ubiquitin ligases: reflections in the magic mirror of cancer. Oncogene, 23, 2028–2036.

18. Shirogane, T., Jin, J., Ang, X.L. and Harper, J.W. (2005) SCFbeta-TRCP controls clock-dependent transcription via casein kinase 1-dependent degradation of the mammalian period-1 (Per1) protein. The Journal of biological chemistry, 280, 26863–26872.

19. Lee, H., Chen, R., Lee, Y., Yoo, S. and Lee, C. (2009) Essential roles of CKIdelta and CKIepsilon in the mammalian circadian clock. Proceedings of the National Academy of Sciences of the United States of America, 106, 21359–21364.

20. Walton, K.M., Fisher, K., Rubitski, D., Marconi, M., Meng, Q.J., Sladek, M., Adams, J., Bass, M., Chandrasekaran, R., Butler, T. et al. (2009) Selective inhibition of casein kinase 1 epsilon minimally alters circadian clock period. J Pharmacol Exp Ther, 330, 430–439.

21. D’Alessandro, M., Beesley, S., Kim, J.K., Jones, Z., Chen, R., Wi, J., Kyle, K., Vera, D., Pagano, M., Nowakowski, R. et al. (2017) Stability of Wake-Sleep Cycles Requires Robust Degradation of the PERIOD Protein. Curr Biol.

22. Gotoh, T., Vila-Caballer, M., Santos, C.S., Liu, J., Yang, J. and Finkielstein, C.V. (2014) The circadian factor Period 2 modulates p53 stability and transcriptional activity in unstressed cells. Molecular biology of the cell.

23. Gotoh, T., Kim, J.K., Liu, J., Vila-Caballer, M., Stauffer, P.E., Tyson, J.J. and Finkielstein, C.V. (2016) Model-driven experimental approach reveals the complex regulatory distribution of p53 by the circadian factor Period 2. Proceedings of the National Academy of Sciences of the United States of America, 113, 13516–13521.

24. Gotoh, T., Vila-Caballer, M., Liu, J., Schiffhauer, S. and Finkielstein, C.V. (2015) Association of the circadian factor Period 2 to p53 influences p53’s function in DNA-damage signaling. Molecular biology of the cell, 26, 359–372.

25. Berndsen, C.E. and Wolberger, C. (2014) New insights into ubiquitin E3 ligase mechanism. Nat Struct Mol Biol, 21, 301–307.

26. Middleton, A.J., Wright, J.D. and Day, C.L. (2017) Regulation of E2s: A Role for Additional Ubiquitin Binding Sites? J Mol Biol, 429, 3430–3440.

27. Balsalobre, A., Damiola, F. and Schibler, U. (1998) A serum shock induces circadian gene expression in mammalian tissue culture cells. Cell, 93, 929–937.

28. Balsalobre, A., Brown, S.A., Marcacci, L., Tronche, F., Kellendonk, C., Reichardt, H.M., Schutz, G. and Schibler, U. (2000) Resetting of circadian time in peripheral tissues by glucocorticoid signaling. Science, 289, 2344–2347.

29. Yang, J., Yan, R., Roy, A., Xu, D., Poisson, J. and Zhang, Y. (2015) The I-TASSER Suite: protein structure and function prediction. Nat Methods, 12, 7–8.

30. Zhang, Y. and Skolnick, J. (2004) SPICKER: a clustering approach to identify near-native protein folds. J Comput Chem, 25, 865–871.

31. Vijay-Kumar, S., Bugg, C.E. and Cook, W.J. (1987) Structure of ubiquitin refined at 1.8 A resolution. J Mol Biol, 194, 531–544.

32. Vanselow, K., Vanselow, J.T., Westermark, P.O., Reischl, S., Maier, B., Korte, T., Herrmann, A., Herzel, H., Schlosser, A. and Kramer, A. (2006) Differential effects of PER2 phosphorylation: molecular basis for the human familial advanced sleep phase syndrome (FASPS). Genes & development, 20, 2660–2672.

33. Vanselow, K. and Kramer, A. (2007) Role of phosphorylation in the mammalian circadian clock. Cold Spring Harb Symp Quant Biol, 72, 167–176.

34. Bunz, F., Dutriaux, A., Lengauer, C., Waldman, T., Zhou, S., Brown, J.P., Sedivy, J.M., Kinzler, K.W. and Vogelstein, B. (1998) Requirement for p53 and p21 to sustain G2 arrest after DNA damage. Science, 282, 1497–1501.

35. Iwakuma, T. and Lozano, G. (2003) MDM2, an introduction. Mol Cancer Res, 1, 993–1000.

36. Honda, R. and Yasuda, H. (2000) Activity of MDM2, a ubiquitin ligase, toward p53 or itself is dependent on the RING finger domain of the ligase. Oncogene, 19, 1473–1476.

37. Chen, J., Marechal, V. and Levine, A.J. (1993) Mapping of the p53 and mdm-2 interaction domains. Mol Cell Biol, 13, 4107–4114.

38. Albrecht, U., Bordon, A., Schmutz, I. and Ripperger, J. (2007) The multiple facets of Per2. Cold Spring Harb Symp Quant Biol, 72, 95–104.

39. Yang, J., Kim, K.D., Lucas, A., Drahos, K.E., Santos, C.S., Mury, S.P., Capelluto, D.G. and Finkielstein, C.V. (2008) A novel heme-regulatory motif mediates heme-dependent degradation of the circadian factor period 2. Mol Cell Biol, 28, 4697–4711.

40. Li, M., Brooks, C.L., Wu-Baer, F., Chen, D., Baer, R. and Gu, W. (2003) Mono-versus polyubiquitination: differential control of p53 fate by Mdm2. Science, 302, 1972–1975.

41. Wenzel, D.M., Stoll, K.E. and Klevit, R.E. (2011) E2s: structurally economical and functionally replete. Biochem J, 433, 31–42.

42. Ye, Y. and Rape, M. (2009) Building ubiquitin chains: E2 enzymes at work. Nat Rev Mol Cell Biol, 10, 755–764.

43. Kruse, J.P. and Gu, W. (2009) Modes of p53 regulation. Cell, 137, 609–622.

44. Toh, K.L., Jones, C.R., He, Y., Eide, E.J., Hinz, W.A., Virshup, D.M., Ptacek, L.J. and Fu, Y.H. (2001) An hPer2 phosphorylation site mutation in familial advanced sleep phase syndrome. Science, 291, 1040–1043.

45. Badura, L., Swanson, T., Adamowicz, W., Adams, J., Cianfrogna, J., Fisher, K., Holland, J., Kleiman, R., Nelson, F., Reynolds, L. et al. (2007) An inhibitor of casein kinase I epsilon induces phase delays in circadian rhythms under free-running and entrained conditions. J Pharmacol Exp Ther, 322, 730–738.

46. Kim, J.K., Forger, D.B., Marconi, M., Wood, D., Doran, A., Wager, T., Chang, C. and Walton, K.M. (2013) Modeling and validating chronic pharmacological manipulation of circadian rhythms. CPT Pharmacometrics Syst Pharmacol, 2, e57.

47. Yoo, S.H., Ko, C.H., Lowrey, P.L., Buhr, E.D., Song, E.J., Chang, S., Yoo, O.J., Yamazaki, S., Lee, C. and Takahashi, J.S. (2005) A noncanonical E-box enhancer drives mouse Period2 circadian oscillations in vivo. Proceedings of the National Academy of Sciences of the United States of America, 102, 2608–2613.

48. Yoo, S.H., Yamazaki, S., Lowrey, P.L., Shimomura, K., Ko, C.H., Buhr, E.D., Siepka, S.M., Hong, H.K., Oh, W.J., Yoo, O.J. et al. (2004) PERIOD2::LUCIFERASE real-time reporting of circadian dynamics reveals persistent circadian oscillations in mouse peripheral tissues. Proceedings of the National Academy of Sciences of the United States of America, 101, 5339–5346.

49. Chen, R., Schirmer, A., Lee, Y., Lee, H., Kumar, V., Yoo, S.H., Takahashi, J.S. and Lee, C. (2009) Rhythmic PER abundance defines a critical nodal point for negative feedback within the circadian clock mechanism. Molecular cell, 36, 417–430.

50. Sasiela, C.A., Stewart, D.H., Kitagaki, J., Safiran, Y.J., Yang, Y., Weissman, A.M., Oberoi, P., Davydov, I.V., Goncharova, E., Beutler, J.A. et al. (2008) Identification of inhibitors for MDM2 ubiquitin ligase activity from natural product extracts by a novel high-throughput electrochemiluminescent screen. J Biomol Screen, 13, 229–237.

51. Clement, J.A., Kitagaki, J., Yang, Y., Saucedo, C.J., O’Keefe, B.R., Weissman, A.M., McKee, T.C. and McMahon, J.B. (2008) Discovery of new pyridoacridine alkaloids from Lissoclinum cf. badium that inhibit the ubiquitin ligase activity of Hdm2 and stabilize p53. Bioorg Med Chem, 16, 10022–10028.

52. Kitagaki, J., Agama, K.K., Pommier, Y., Yang, Y. and Weissman, A.M. (2008) Targeting tumor cells expressing p53 with a water-soluble inhibitor of Hdm2. Mol Cancer Ther, 7, 2445–2454.

53. Dickens, M.P., Fitzgerald, R. and Fischer, P.M. (2010) Small-molecule inhibitors of MDM2 as new anticancer therapeutics. Semin Cancer Biol, 20, 10–18.

54. Schwartz, A.L. and Ciechanover, A. (2009) Targeting proteins for destruction by the ubiquitin system: implications for human pathobiology. Annu Rev Pharmacol Toxicol, 49, 73–96.

55. Stojkovic, K., Wing, S.S. and Cermakian, N. (2014) A central role for ubiquitination within a circadian clock protein modification code. Front Mol Neurosci, 7, 69.

56. Smolensky, M.H., Hermida, R.C., Reinberg, A., Sackett-Lundeen, L. and Portaluppi, F. (2016) Circadian disruption: New clinical perspective of disease pathology and basis for chronotherapeutic intervention. Chronobiol Int, 33, 1101–1119.

57. Ohsaki, K., Oishi, K., Kozono, Y., Nakayama, K., Nakayama, K.I. and Ishida, N. (2008) The role of {beta}-TrCP1 and {beta}-TrCP2 in circadian rhythm generation by mediating degradation of clock protein PER2. J Biochem, 144, 609–618.

58. Lamia, K.A., Sachdeva, U.M., DiTacchio, L., Williams, E.C., Alvarez, J.G., Egan, D.F., Vasquez, D.S., Juguilon, H., Panda, S., Shaw, R.J. et al. (2009) AMPK regulates the circadian clock by cryptochrome phosphorylation and degradation. Science, 326, 437–440.

59. Zhao, X., Hirota, T., Han, X., Cho, H., Chong, L.W., Lamia, K., Liu, S., Atkins, A.R., Banayo, E., Liddle, C. et al. (2016) Circadian Amplitude Regulation via FBXW7-Targeted REV-ERBalpha Degradation. Cell, 165, 1644–1657.

60. Sahar, S., Zocchi, L., Kinoshita, C., Borrelli, E. and Sassone-Corsi, P. (2010) Regulation of BMAL1 protein stability and circadian function by GSK3beta-mediated phosphorylation. PLoS One, 5, e8561.

61. Ruffner, H., Bauer, A. and Bouwmeester, T. (2007) Human protein-protein interaction networks and the value for drug discovery. Drug Discov Today, 12, 709–716.

62. DeBruyne, J.P., Baggs, J.E., Sato, T.K. and Hogenesch, J.B. (2015) Ubiquitin ligase Siah2 regulates RevErbalpha degradation and the mammalian circadian clock. Proceedings of the National Academy of Sciences of the United States of America, 112, 12420–12425.

63. Brenkman, A.B., de Keizer, P.L., van den Broek, N.J., Jochemsen, A.G. and Burgering, B.M. (2008) Mdm2 induces mono-ubiquitination of FOXO4. PLoS One, 3, e2819.

64. Marine, J.C. and Lozano, G. (2010) Mdm2-mediated ubiquitylation: p53 and beyond. Cell death and differentiation, 17, 93–102.

65. Grossman, S.R., Deato, M.E., Brignone, C., Chan, H.M., Kung, A.L., Tagami, H., Nakatani, Y. and Livingston, D.M. (2003) Polyubiquitination of p53 by a ubiquitin ligase activity of p300. Science, 300, 342–344.

66. Lai, Z., Ferry, K.V., Diamond, M.A., Wee, K.E., Kim, Y.B., Ma, J., Yang, T., Benfield, P.A., Copeland, R.A. and Auger, K.R. (2001) Human mdm2 mediates multiple mono-ubiquitination of p53 by a mechanism requiring enzyme isomerization. The Journal of biological chemistry, 276, 31357–31367.

67. Oklejewicz, M., Destici, E., Tamanini, F., Hut, R.A., Janssens, R. and van der Horst, G.T. (2008) Phase resetting of the mammalian circadian clock by DNA damage. Curr Biol, 18, 286–291.

68. Papp, S.J., Huber, A.L., Jordan, S.D., Kriebs, A., Nguyen, M., Moresco, J.J., Yates, J.R. and Lamia, K.A. (2015) DNA damage shifts circadian clock time via Hausp-dependent Cry1 stabilization. Elife, 4.

69. Gaddameedhi, S., Reardon, J.T., Ye, R., Ozturk, N. and Sancar, A. (2012) Effect of circadian clock mutations on DNA damage response in mammalian cells. Cell cycle, 11, 3481–3491.

